# Comparing 3D genome organization in multiple species using Phylo-HMRF

**DOI:** 10.1101/552505

**Authors:** Yang Yang, Yang Zhang, Bing Ren, Jesse Dixon, Jian Ma

## Abstract

Recent developments in whole-genome mapping approaches for the chromatin interactome (such as Hi-C) have offered new insights into 3D genome organization. However, our knowledge of the evolutionary patterns of 3D genome structures in mammalian species remains surprisingly limited. In particular, there are no existing phylogenetic-model based methods to analyze chromatin interactions as continuous features across different species. Here we develop a new probabilistic model, named phylogenetic hidden Markov random field (Phylo-HMRF), to identify evolutionary patterns of 3D genome structures based on multi-species Hi-C data by jointly utilizing spatial constraints among genomic loci and continuous-trait evolutionary models. The effectiveness of Phylo-HMRF is demonstrated in both simulation evaluation and application to real Hi-C data. We used Phylo-HMRF to uncover cross-species 3D genome patterns based on Hi-C data from the same cell type in four primate species (human, chimpanzee, bonobo, and gorilla). The identified evolutionary patterns of 3D genome organization correlate with features of genome structure and function, including long-range interactions, topologically-associating domains (TADs), and replication timing patterns. This work provides a new framework that utilizes general types of spatial constraints to identify evolutionary patterns of continuous genomic features and has the potential to reveal the evolutionary principles of 3D genome organization.

## Introduction

In humans and other higher eukaryotes, chromosomes are folded and organized in three-dimensional (3D) space and different chromosomal loci interact with each other (Bonev and Cavalli, 2016; Rowley and Corces, 2018). Recent developments in whole-genome mapping approaches for the chromatin interactome such as Hi-C (Lieberman-Aiden et al., 2009; Rao et al., 2014) and ChIA-PET (Tang et al., 2015) have facilitated the identification of genome-wide chromatin organizations comprehensively, revealing important 3D genome features such as loops (Rao et al., 2014; Tang et al., 2015), topologically-associating domains (TADs) (Dixon et al., 2012; Nora et al., 2012), and A/B compartments (Lieberman-Aiden et al., 2009). A limited number of attempts have been made to analyze these 3D genome features across different species. An earlier study using Hi-C showed that the positions of TADs were largely conserved between human and mouse within syntenic genomic regions (Dixon et al., 2012). Another study demonstrated that evolutionary changes in TAD structure correspond with the creation or elimination of CTCF binding sites using relatively low resolution Hi-C data from rhesus macaque, dog, rabbit, and mouse (Rudan et al., 2015). More recently, TADs have been shown to have strong conservation in mammalian evolution with the TADs boundaries under potential negative selections against genome rearrangements (Lazar et al., 2018; Fudenberg and Pollard, 2019). These previous analyses pointed to the conservation and changes of 3D genome structure across different species, although a more comprehensive characterization of the detailed evolutionary patterns of 3D genome organization remains unclear. Additionally, as most of the initial comparative analysis of 3D genomes focused primarily on distantly related organisms, there is limited understanding of how 3D genome features may have evolved in closely related mammalian species, especially in recent primate evolution which is of particular interest to understand human specific and great-ape specific gene regulations.

On the algorithmic side, existing computational approaches for comparing 3D genome organization across multiple species have surprisingly limited capability. Importantly, multi-species functional genomic data from various high-throughput epigenomic assays (e.g., Hi-C, ChIP-seq, Repli-seq) are continuous traits in nature. However, such continuous signals are often converted to discrete values to identify distinctive feature patterns (e.g., presence or absence of TADs) for subsequent comparisons, which may cause dramatic loss of information of more subtle differences from the original data. Although methods have been developed to quantitatively analyze the strengths of chromatin interactions from Hi-C data, to the best of our knowledge, there are no existing phylogenetic-model based methods available to analyze Hi-C data as continuous signals across different species in a genome-wide manner to uncover evolutionary patterns of 3D genome organization.

We previously developed a method called phylogenetic hidden Markov Gaussian Processes (Phylo-HMGP) (Yang et al., 2018) to estimate evolutionary patterns given continuous functional genomic data (e.g., Repli-seq) along the genome from multiple species. Phylo-HMGP considers evolutionary affinities among species in a hidden Markov model (HMM), utilizing evolutionary constraints and also dependencies along one-dimensional (1D) genome coordinates. However, the HMMs, as used by Phylo-HMGP, are based on 1D Markov chains, which cannot be simply used to model generalized spatial dependencies (such as those reflected in Hi-C data) to consider the interactions between nodes in an arbitrary graph. Therefore, HMM-based methods cannot be directly applied to discovering patterns of higher-order chromatin interactions from Hi-C contact matrices, which consist of continuous measurements of contact frequencies between a pair of genomic loci.

Here, we develop a new probabilistic model, phylogenetic hidden Markov random field (Phylo-HMRF), which integrates the continuous-trait evolutionary constraints with the hidden Markov random field (HMRF) model, to capture evolutionary patterns of continuous genomic features across species by utilizing generalized spatial constraints. We demonstrated the advantage of Phylo-HMRF using simulation data. In addition, we applied Phylo-HMRF to a new Hi-C dataset from the same cell type (lymphoblastoid cell) in four primate species (human, chimpanzee, bonobo, and gorilla). Phylo-HMRF identified different evolutionary patterns of Hi-C contacts across the four species, including both conserved patterns and lineage-specific patterns. These patterns show strong correlations with long-range Hi-C interactions, TAD structures, and the evolutionary patterns of DNA replication timing in primate species. Phylo-HMRF offers an effective model to potentially reveal important evolutionary principles of 3D genome organization. The source code of Phylo-HMRF can be accessed at: https://github.com/macompbio/Phylo-HMRF.

## Results

### Overview of the Phylo-HMRF model

The overview of the Phylo-HMRF method is shown in Fig. 1. Our goal is to identify different evolutionary patterns of chromatin conformation from multi-species Hi-C data. The input contains the Hi-C contact frequency data from each species. We align the Hi-C contacts in each species to the reference genome, and obtain a combined multi-species Hi-C contact map based on the reference genome as shown in Fig. 1A, where each node in the map corresponds to multi-species contact frequencies between the corresponding pair of genomic loci (Supplementary Methods). Hence each node is associated with a multi-dimensional feature as the multi-species observation. We also assume that each node has a hidden state that represents the evolutionary pattern of Hi-C contacts between the corresponding pair of genomic loci. Phylo-HMRF estimates the hidden state of each node by considering both spatial dependencies among nodes encoded by an HMRF and the evolutionary dependencies between species in the phylogeny. The continuous-trait evolutionary models are embedded into the HMRF. Therefore, each hidden state corresponds to an evolutionary model that is represented by a parameterized phylogenetic tree. The output of Phylo-HMRF contains the partition of the combined multi-species Hi-C contact map, where adjacent nodes with the same hidden state are in the same partition. These partitions reflect the distribution of different evolutionary patterns of Hi-C contact frequencies. As shown in Fig. 1B, Phylo-HMRF uses the Ornstein-Uhlenbeck (OU) process as the continuous-trait evolutionary model. Fig. 1C is an illustration of the possible Hi-C evolutionary patterns that Phylo-HMRF aims to uncover across four primate species in this work.

**Figure 1:**
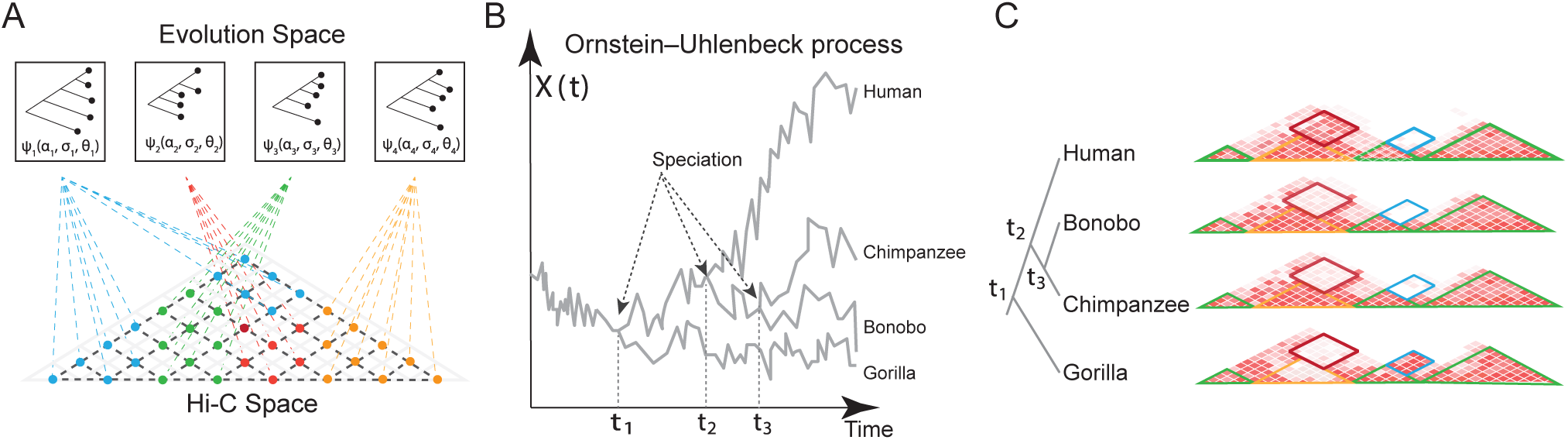
Overview of Phylo-HMRF. **(A)** Illustration of the possible evolutionary patterns of chromatin interaction. The Hi-C space is a combined multi-species Hi-C contact map, which integrates aligned Hi-C contact maps of each species. Each node represents the multi-species observations of Hi-C contact frequency between a pair of genomic loci, with a hidden state assigned. Nodes with the same color have the same hidden state and are associated with the same type of evolutionary pattern represented by a parameterized phylogenetic tree *ψ*_*i*_. The parameters of *ψ*_*i*_ include the selection strengths *α*_*i*_, Brownian motion intensities *σ*_*i*_, and the optimal values *θ*_*i*_ based on the Ornstein-Uhlenback (OU) process assumption. **(B)** Illustration of the OU process over a phylogenetic tree with four observed species. Time axis represents the evolution history. *X*(*t*) represents the trait at time *t*. The trajectories reflect the evolution of the continuous-trait features in different lineages, where the time points *t*_1_, *t*_2_, *t*_3_ represent the speciation events. **(C)** A cartoon example of the possible evolutionary patterns (partitioned with different colors). Phylo-HMRF aims to identify evolutionary Hi-C contact patterns among four primate species in this work. The four Hi-C contact maps represent the observations from the four species, which are combined into one multi-species Hi-C map as the input to Phylo-HMRF, as shown in **(A)**. The phylogenetic tree of the four species in this study is on the left. The partitions with green borders are conserved Hi-C contact patterns. The partitions with red or blue borders represent lineage-specific Hi-C contact patterns.

Note that an evolutionary pattern identified by Phylo-HMRF is associated with the conservation or variation of the feature of interest across different species. For example, in this study, we may observe conserved high Hi-C contacts in all the compared species between a specific pair of genomic loci. We may also observe that strong Hi-C contacts only exist in some of the species between a specific pair of genomic loci. These different types of feature distribution across species are representative of the evolutionary patterns that we want to identify as states. In addition, Phylo-HMRF provides a framework to utilize both general types of spatial dependencies among genomic loci and evolutionary relationships among species to identify evolutionary patterns from multi-species continuous-trait features. The general types of spatial dependencies refer to any type of dependencies that can be represented by the edge connections in a graph.

### Performance evaluation of Phylo-HMRF using simulation

We evaluated the performance of Phylo-HMRF using simulations to demonstrate improvement in identifying 2D evolutionary feature patterns. We applied Phylo-HMRF to 16 simulated datasets in two sets of simulations, each of which contains 8 datasets. Suppose the samples in simulated datasets correspond to nodes in a graph ***𝒢***. Similar to a Hi-C contact map, ***𝒢*** has a 2D lattice structure of size *n × n*, where each vertex is associated with a sample. The samples thus represent features of vertices in ***𝒢***. For example, a sample can represent the interaction intensities between the *i*-th locus and the *j*-th locus out of *n* genomic loci in multiple species, (1 *≤ i, j ≤ n*). We also assume that each sample has a class label, or hidden state. Each hidden state is associated with an emission probability function and determines the observed feature of the sample with this state. The hidden states of the samples are assumed to be from a Markov random field (MRF). Thus, the hidden state of a sample is spatially dependent on the hidden states of its neighbors in ***𝒢***. In the simulation evaluation, we first simulated the hidden state configurations of the samples by simulating an MRF through Gibbs sampling (Geman and Geman, 1984). We then simulated the multi-species observations from multi-variate Gaussian distributions with OU model parameters embedded. The details of the data simulation are in Supplementary Methods.

Based on the simulated datasets, we compared Phylo-HMRF with several other methods, including the Gaussian-HMRF method (Zhang et al., 2001), the Gaussian mixture model (GMM), and the *K*-means clustering method, to infer hidden states of the samples. Each method was run repeatedly for 10 times on each simulated dataset with different random initialization, given the number of hidden states *M* = 10. Additionally, we also included two image segmentation methods SLIC (Simple Linear Iterative Clustering) (Achanta et al., 2012) and Quick Shift (Vedaldi and Soatto, 2008) for comparison (Supplementary Methods). Both methods have been effectively used in image segmentation applications. The two methods were also run repeatedly for 10 times each. We adjusted the parameter configurations of each of the two methods such that the number of output segments is 10. To utilize the image segmentation methods for state estimation across species, we considered the graph ***𝒢*** as an image and considered the features of each species as one color channel of the image. The average performance of the 10 results for each method was reported as the final performance of the corresponding method with respect to different types of evaluation metrics. We evaluated the performance of each method by comparing the predicted states and ground truth hidden states, using evaluation metrics Normalized Mutual Information (NMI), Adjusted Mutual Information (AMI), Adjusted Rand Index (ARI), Precision, Recall, and *F*_1_ score (Manning et al., 2008; Vinh et al., 2010) (Supplementary Methods).

The evaluation results are shown in Fig. 2 and Table S1. We found that Phylo-HMRF outforms all the other methods on different types of evaluation metrics in each simulated dataset in simulation study I. Phylo-HMRF consistently outperforms Gaussian-HMRF, demonstrating that encoding the evolution information can improve accuracy. Even though all the multi-species observations are simulated from Gaussian distribution, using Gaussian distribution alone in inference may not reveal the possible evolutionary dependencies between the species. In addition, both Phylo-HMRF and Gaussian-HMRF show advantage over GMM and *K*-means clustering, suggesting that encoding the spatial constraints is also crucial. Moreover, Phylo-HMRF outperforms the two image segmentation methods SLIC and Quick Shift in different simulated datasets. The image segmentation methods perform segmentation of the image representation of the cross-species data based on feature similarity and spatial proximity. Regions that belong to the same evolutionary pattern (e.g., conserved high in Hi-C contacts across species) can be assigned different labels if they are distant from each other in spatial location, which affects the accuracy and interpretability of hidden state estimation.

**Figure 2:**
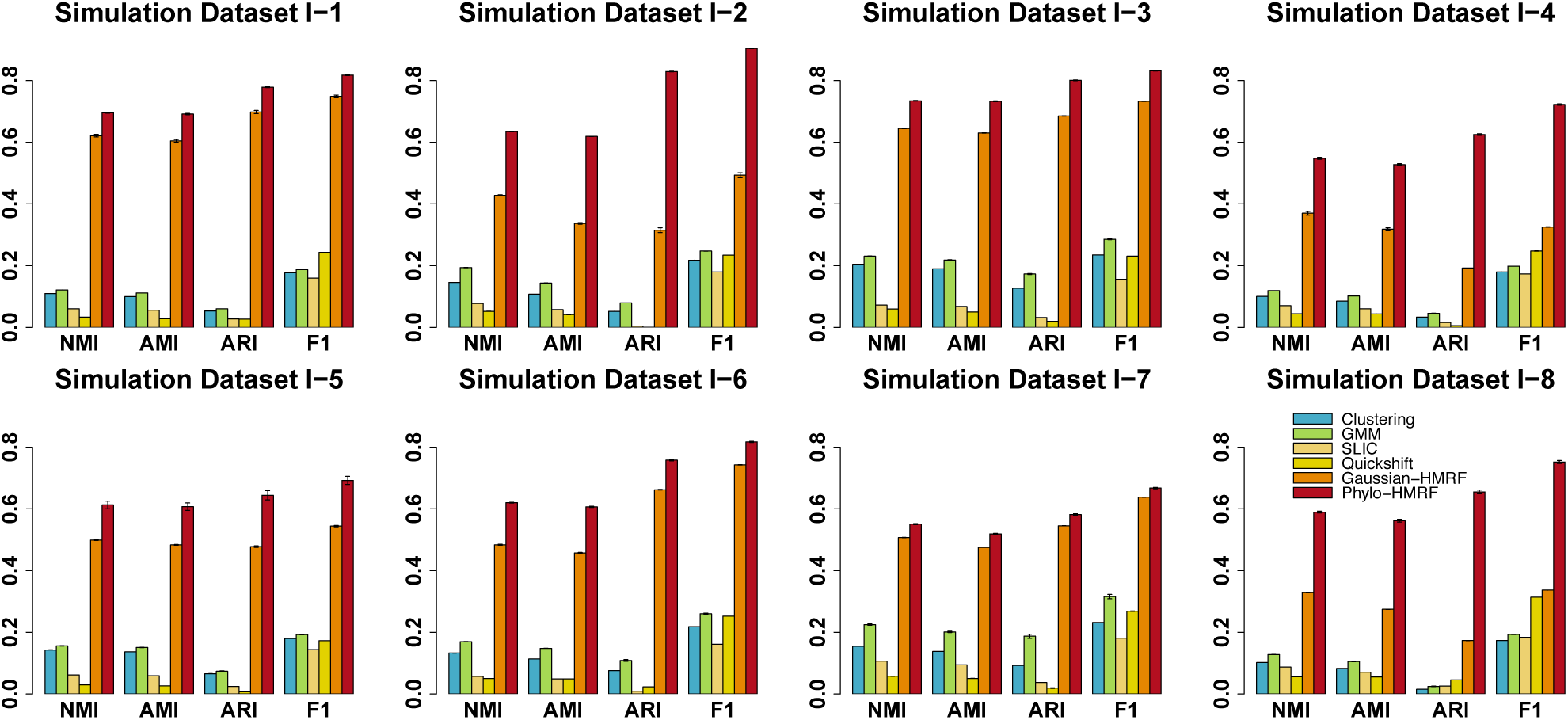
Performance evaluation of *K*-means Clustering, GMM, SLIC, Quick Shift, Gaussian-HMRF, and Phylo-HMRF on eight simulation datasets in simulation study I with respect to NMI (Normalized Mutual Information), AMI (Adjusted Mutual Information), ARI (Adjusted Rand Index), and *F*_1_ score. The standard error of the results from 10 runs of each method is shown as the error bar, respectively.

We further performed simulation study II, where we simulated another eight datasets with different parameter configurations, in order to assess if the advantage of Phylo-HMRF is consistent over varied simulation parameter settings (Supplementary Methods). The eight sets of simulated hidden states are shared between simulation studies I and II, while the observable random fields are simulated with different parameter settings. We then applied Phylo-HMRF and the other methods to the datasets in simulation study II and evaluated performance using the same procedure as we used in simutation study I. The evaluation results are shown in Fig. S1. Again we found that Phylo-HMRF consistently outperforms the other methods across all datasets in simulation study II. Taken together, the simulation evaluation demonstrated that Phylo-HMRF is able to achieve improved accuracy consistently in estimating evolutionary patterns of Hi-C contacts in multiple species.

### Phylo-HMRF identifies different Hi-C contact patterns across multiple primate species

We applied Phylo-HMRF to a Hi-C dataset from four primate species. We used the Hi-C data in GM12878 in human from the 4DN data portal. We generated Hi-C data from the lymphoblastoid cells of three other primate species, including chimpanzee, bonobo, and gorilla. There are 290M, 270M, 240M, and 290M mapped read pairs for the four primate species, respectively. The genome assemblies used for the four species are hg38, panTro5, panPan2, and gorGor4, respectively.

We ran Phylo-HMRF on all the syntenic regions on autosomes based on the human genome. We first identified 92 synteny blocks in 50 kb resolution (i.e., ignoring rearrangements smaller than 50 kb among the four species) using the method inferCARs (Ma et al., 2006), covering 92.64% of the sequenced regions in the human genome (Fig. S2, Supplementary Methods). For example, we identified 9 major synteny blocks on human chromosome 1 among the four species, covering 92.50% of human chromosome 1 (Fig. 3B). We then applied Phylo-HMRF to perform genome-wide evolutionary pattern estimation over the multiple synteny blocks across different chromosomes jointly, with each synteny block represented as a subgraph of ***𝒢***. The number of states is set to be 30 based on estimation from the results of *K*-means clustering (Supplementary Methods, Fig. S3).

**Figure 3:**
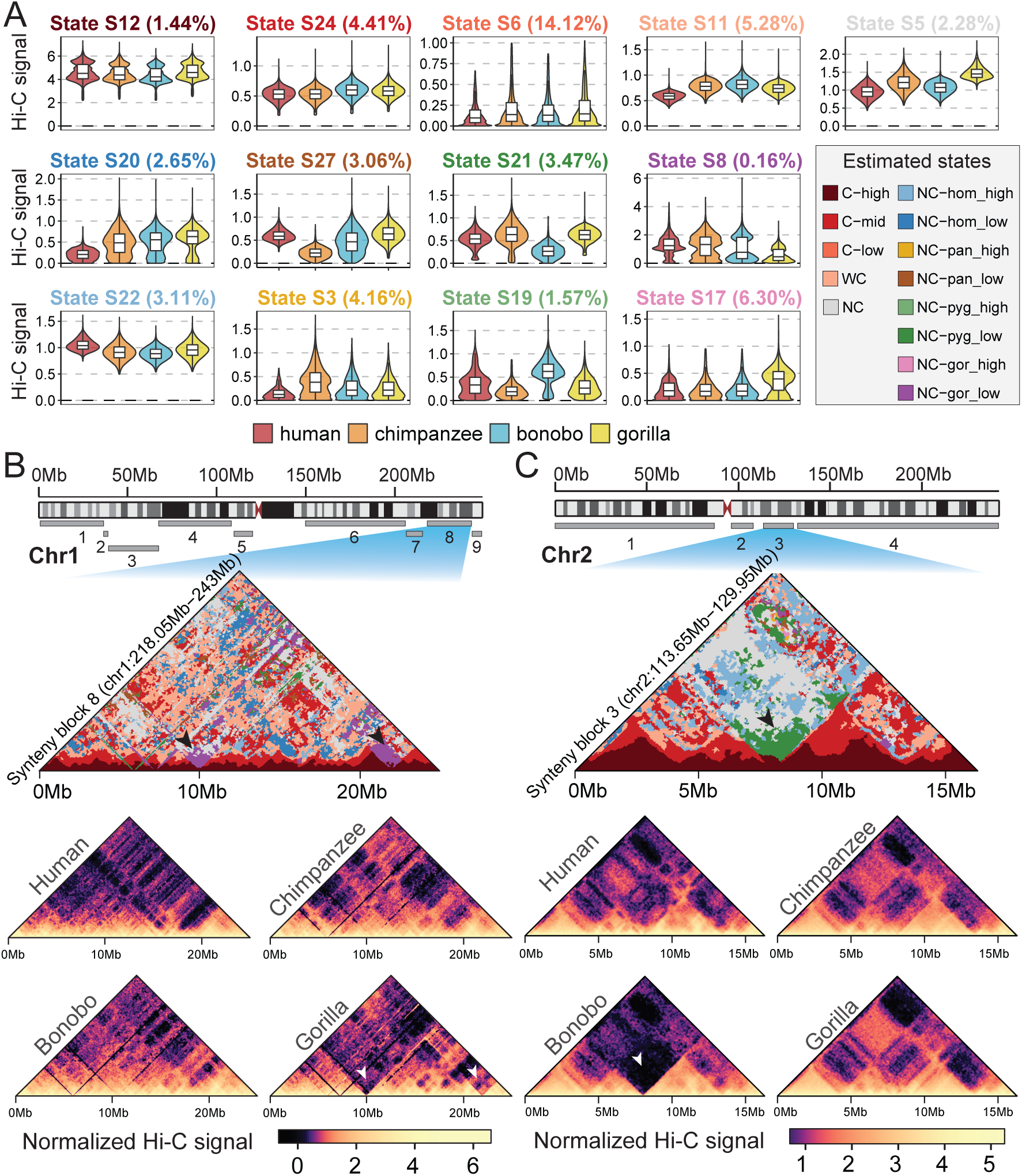
Evolutionary patterns of Hi-C contact frequency estimated by Phylo-HMRF. **(A)** Representative states from the 13 groups of evolutionary patterns. One state from each group is presented. The boxplots show the normalized cross-species Hi-C contact frequency distributions of the four species in the corresponding states with outliers removed. **(B)** Cross-species Hi-C contact frequency states identified in synteny block 8 on chromosome 1, in comparison with the Hi-C contact maps of the four primate species. First row: Locations of the nine identified synteny blocks on chromosome 1. Second row: Cross-species Hi-C contact frequency states identified by Phylo-HMRF. The black arrows point to two examples of identified gorilla-specific low Hi-C contact frequency state (NC-gor low, purple color) in the combined Hi-C contact map. Third and fourth rows: Hi-C contact maps of the four primate species in this synteny block, with signal scale displayed at the bottom. Darker color in the Hi-C contact map represents lower contact frequency. The white arrows point to the corresponding locations of the two examples of identified gorilla-specific low contact state. **(C)** Cross-species Hi-C contact frequency states identified in synteny block 3 on chromosome 2. First row: Locations of the four identified synteny blocks on chromosome 2. Second row: Cross-species Hi-C contact frequency states identified by Phylo-HMRF. The black arrow points to one example of identified bonobo-specific low Hi-C contact state (NC-pyg low, green color) in the combined Hi-C contact map. The white arrow points to the corresponding location of the example of identified bonobo-specific low state in the Hi-C contact maps of the four species.

Phylo-HMRF identified both conserved and lineage-specific evolutionary patterns of Hi-C contact frequencies across the four primate species. We further categorized the 30 estimated hidden states into 13 groups which show higher-level distinctiveness of heterogeneous evolutionary patterns (Supplementary Methods). Four of the groups represent conserved or weakly conserved cross-species patterns in Hi-C contact frequency, which are conserved high in Hi-C contact frequency (C-high), conserved middle-level (C-mid), conserved low (C-low), and weakly conserved middle-level (WC). The four groups cover 51.14% of all the nodes in the cross-species Hi-C maps of the synteny blocks. Nine of the groups correspond to non-conserved evolutionary patterns in Hi-C contacts, where eight exhibit lineage-specific patterns. Specifically, the nine groups are human-specific high in Hi-C contact frequency (NC-hom high), human-specific low (NC-hom low), chimpanzee-specific high (NC-pan high), chimpanzee-specific low (NC-pan low), bonobo-specific high (NC-pyg high), bonobo-specific low (NC-pyg low), gorilla-specific high (NC-gor high), gorilla-specific low (NC-gor low) and non-conserved (NC). Examples of the representative estimated states from each of the 13 groups are shown in Fig. 3A. Hi-C contact frequency distributions of multiple species in all 30 estimated states are shown in Fig. S4.

In Fig. 3B and Fig. 3C, as examples we show the estimated states in synteny block 8 on chromosome 1 and in synteny block 3 on chromosome 2, along with the input Hi-C contact maps of the four species. The rotated upper triangular matrix as shown in the second row of Fig. 3B or Fig. 3C represents the estimated hidden states of the graph ***𝒢*** of the HMRF in Hi-C data comparison in the corresponding synteny block, which is of the same size as the multi-species Hi-C contact maps in this synteny block. Each vertex in ***𝒢*** corresponds to a pair of genomic loci in the Hi-C contact map. The hidden state configuration is visualized as an image, where different colors represent different estimated hidden states. Adjacent nodes that are assigned to the same hidden state form a contiguous segment in the image. Therefore, based on the estimated states, ***𝒢*** is partitioned, reflecting different cross-species Hi-C contact patterns. In Fig. 3B, we found that there are two gorilla-specific low Hi-C contact patterns near the diagonal area, which are colored in purple. These two regions correspond to the gorilla-specific low Hi-C states that appear in the state-distance plots of synteny block 8 on chromosome 1 in Fig. S5. In Fig. 3C, we observed that there is a bonobo-specific low Hi-C contact frequency pattern detected near the diagonal area, which is colored in green. By comparing the estimated hidden states to the corresponding Hi-C contact maps of the four species, we found that the estimated states accurately reflect what can be observed in Hi-C contact maps in different species.

Next we compared the distributions of evolutionary patterns of Hi-C contacts over changing distances between a pair of genomic loci in each synteny block. We consider that Hi-C contacts over short genomic loci distances are local Hi-C contacts (genomic loci distance *<*3Mb), and Hi-C contacts over large distances represent longer-range contacts. Short genomic loci distances correspond to an area near the diagonal of the Hi-C contact map. The state-distance plots across the synteny blocks on all the autosomes are shown in Fig. 4A. We observed that C-high, C-mid, and WC states are the predominant states around the diagonal area when the genomic loci distance is within the range of around 2.5 Mb (black arrow in Fig. 4A). This suggests that the majority of the local Hi-C contact patterns and the associated genome structures are likely to be conserved across difference species. At large genomic loci distances, which corresponds to the off-diagonal area in the Hi-C contact map, the C-low and non-conserved states have much larger percentages as expected. We also found that the distribution over genomic loci distance varies across different lineage-specific states. For single synteny blocks, the state-distance plots of the major synteny blocks on chromosome 1 and chromosome 2 are shown in Fig. S5 and Fig. S6 as examples. Overall, we found that there are similar enrichment patterns of states within a short distance range across the synteny blocks, while different blocks also exhibit varied trends of how evolutionary patterns are distributed at different genomic loci distances. We also observed that the lineage-specific states are distributed unevenly among the synteny blocks, showing occurrences either in local Hi-C contacts or long-range Hi-C contacts.

**Figure 4:**
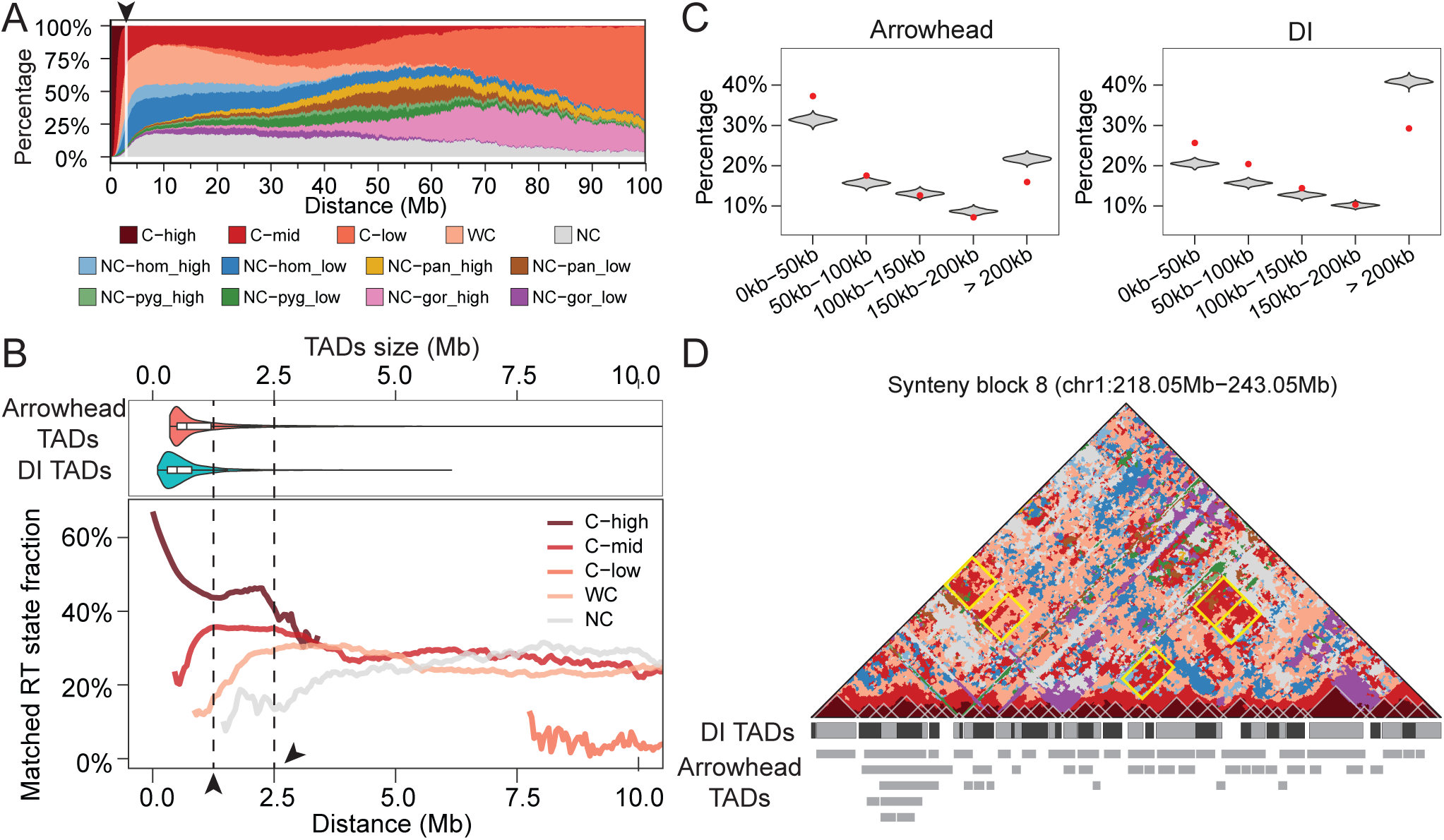
Comparison between the evolutionary states of Hi-C contacts estimated by Phylo-HMRF and other features of genome structure and function. **(A)** Global state-distance plots show the enrichment of different evolutionary patterns at different distances between genomic loci across synteny blocks on all autosomes. The black arrow points to the distance range around 2.5Mb where C-high and C-mid are the predominant states. **(B)** The percentage of paired genomic loci that is conserved in RT in five Hi-C contact evolutionary pattern groups. The top panel shows the distributions of TAD sizes, aligned with the axis of the genomic loci distance. The first black arrow points to the position of the first observed changing point on the Matched-RT state fraction curve for the C-high state and for the C-mid state. The second black arrow points to the position of 2.5MB, where the trend change is observed in the state-distance plot as shown in **(A)**. **(C)** Distributions of distance between the boundaries of all the identified local conserved high Hi-C contact patterns and the nearest TAD boundaries for both Arrowhead TADs and DI TADs. **(D)** Comparison between the boundaries of local conserved high Hi-C contact patterns and the TAD boundaries in synteny block 8 on chromosome 1. Several potential long-range conserved TAD interaction patterns are shown with yellow solid lines as borders.

Together these results demonstrate the effectiveness of Phylo-HMRF to identify evolutionary patterns of Hi-C contacts across different species in a phylogeny.

### Hi-C evolutionary patterns correlate with DNA replication timing patterns

We next compared the predicted states from Phylo-HMRF with other features of genome structure and function. We previously reported evolutionary patterns of DNA replication timing (RT) using Repli-seq of multiple primate species (Yang et al., 2018). We identified 5 groups of RT evolutionary states, which are conserved early in RT (E), weakly conserved early (WE), conserved late (L), weakly conserved late (WL), and non-conserved (NC). For each pair of genomic loci with estimated Hi-C states, we examined the RT state composition of the corresponding two genomic loci. If the paired genomic loci share similar conserved RT states, i.e., both are E/WE or both are L/WL, we annotated this pair as conserved in RT (C), otherwise we annotated it as non-conserved in RT. We then computed the percentage of contact loci that have the conserved RT states in the C-high, C-mid, WC, C-low, and NC Hi-C states identified by Phylo-HMRF over a range of different distances (0-10Mb). The results are shown in Fig. 4B. Notably, it is clear that C-high, C-mid, and WC Hi-C contacts patterns have higher enrichment of genome contacts with conserved RT patterns than the NC group over most of the distance ranges. The percentage is particularly high in the C-high state for genomic loci that are less than 4 Mb apart. This suggests that the conserved high Hi-C contact states are strongly correlated with those genomic loci pairs that both have consistent conserved RT patterns across species. We further explored potential connections between the features and the curves observed in Fig. 4B with known chromatin structure patterns such as TADs. We considered the TADs in GM12878 in human called using the Arrowhead method (Rao et al., 2014) and the Directionality Index (DI) method (Dixon et al., 2012), which are named Arrowhead TAD and DI TAD, respectively. The average sizes of Arrowhead TADs and DI TADs are around 1Mb and 0.6Mb, respectively. As shown in Fig. 4B, there is a changing point around 1MB in the distance axis on the Matched-RT state fraction curve both for the C-high state and for the C-mid state, which approximately matches the average size of TADs. This implies that the identified evolutionary patterns of local high Hi-C contacts and the evolutionary patterns of RT states may be constrained by the TADs structures.

### Hi-C evolutionary patterns correlate with TADs

TADs are important higher-order genome organization features revealed by Hi-C (Dixon et al., 2012). Here we focus on the diagonal of the Hi-C contact maps that reflect local Hi-C contact patterns across species. For segments of estimated Hi-C states on the diagonal, we used windows that could match the segments to detect the block patterns (Supplementary Methods). We identified 2,793 block patterns on the diagonals of the Hi-C contact maps of all the synteny blocks on all chromosomes.

We compared the boundaries of the diagonal blocks detected from the states predicted by Phylo-HMRF with TADs boundaries called using Arrowhead and DI, respectively. For each boundary of every identified diagonal block, we calculated the distance between the block boundary and the nearest TAD boundary. We then computed the percentages of the diagonal blocks whose distance to the nearest TAD boundaries fall in different distance ranges. Since the Hi-C contact frequency was measured at a resolution of 50Kb in this study, the boundaries of the diagonal blocks detected from the estimated states are all at 50Kb resolution. The distance between a diagonal block boundary and a nearest TAD boundary is calculated in increments of 50Kb (one genome bin) accordingly. We also estimated the empirical distributions of the distances between boundaries of a possible diagonal block and a nearest TAD by randomly shuffling the identified diagonal blocks (Supplementary Methods).

We observed that the distance between the boundaries of the identified diagonal blocks and the nearest TADs are significantly more enriched in the distance intervals that represent relatively small distances (e.g., [0,50Kb] and (50Kb,100Kb]) than expected (Fig. 4C). Specifically, 55.60% of the identified diagonal block boundaries are matched by an Arrowhead TAD boundary with the distance less than 2 bins, which is significantly higher than the corresponding expected percentage 36.08% observed from the empirical distribution (empirical *p*-value*<*2e-03). Similarly, the percentages of the identified diagonal block boundaries that are matched by a DI TAD boundary within 1 bin or 2 bins are significantly higher than the expected percentages (empirical *p*-value*<*1e-03). In contrast, the percentage of the diagonal block boundaries with distance to a nearest TAD boundary larger than 4 bins are significantly smaller than expected (empirical *p*-value*<*1e-03).

For example, we identified 34 blocks along the diagonal of the Hi-C map of synteny block 8 on chromosome 1 from the estimated Hi-C evolutionary states based on Phylo-HMRF. The identified blocks are all C-high states, which are shown with white borders in Fig. 4D. We observed that the block boundaries show high consistency with the TAD boundaries (Fig. 4D). Specifically, 70.77% of the block boundaries match an Arrowhead TAD boundary or a DI TAD boundary within 2 bins. The capability of Phylo-HMRF in detecting TAD boundaries without using a TAD calling algorithm implies that TADs are an important type of units of genome organization evolution. The result also reflects the accuracy of Phylo-HMRF in estimating Hi-C evolutionary patterns.

Furthermore, we observed C-mid and WC states in off-diagonal area of the Hi-C contact map, which potentially correspond to the long-range interactions between two TADs that are conserved across species. Five examples of the potentially conserved long-range TADs interactions are shown in Fig. 4D (highlighted in yellow borders).

Taken together, our results suggest that TADs are important 3D genome organization features in genome evolution in primate species. In addition, the evolutionary changes of intra-TAD interactions (i.e., local contacts) and inter-TAD interactions (i.e., long-range contacts) can be uncovered effectively by Phylo-HMRF.

## Discussion

In this work, we developed Phylo-HMRF, a continuous-trait probabilistic model that provides a new framework to utilize spatial dependencies among genomic loci in 3D space to identify evolutionary patterns of Hi-C contacts across different species in a phylogeny. We applied Phylo-HMRF to analyze Hi-C data from the lymphoblastoid cells in four primate species (human, chimpanzee, bonobo, and gorilla). Phylo-HMRF is able to identify different genome-wide cross-species Hi-C contact patterns, including conserved and lineage-specific patterns in both local interactions and long-range interactions. The identified evolutionary patterns of 3D genome structure have strong correlation with other types of genomic structural and functional features such as TADs and DNA replication timing. From a methodology standpoint, Phylo-HMRF is a flexible framework that can also be applied to other types of multi-species continuous-trait features where there are 2D or 3D spatial dependencies for the features among the genomic loci. Overall, through a proof-of-principle application, we demonstrate that Phylo-HMRF is a promising new method to effectively uncover detailed evolutionary patterns of 3D genome organization based on Hi-C across multiple species.

There are several aspects where our method can be improved. First, model selection methods such as the utilization of the AIC and BIC criterion (Akaike, 1973; Schwarz et al., 1978) may help select the number of states more automatically. Second, Phylo-HMRF has only been applied to synteny blocks across species at the moment and does not explicitly model the chromatin conformation differences due to large-scale genome rearrangements in evolution. It will be an important next step to model genome rearrangements and genome organization evolution in an integrative manner. Third, to study a larger number of more distantly related species, we may face several challenges. As the number of model parameters and feature dimensions increase linearly with the tree size, both the computation demand increases and the model is exposed to a higher possibility of local minima and overfitting especially for a smaller sample size of multi-species feature observations. There will also be more potential misalignments among genomes of distantly related species, resulting in fewer available samples. It will be useful to incorporate more efficient parameter regularization which is compatible with the evolutionary models in the optimization part of Phylo-HMRF, and to develop imputation methods for the missing observations in multi-species genomic data, especially in large-scale phylogenetic trees. Fourth, we assume that all the phylogenetic trees associated with different hidden states share the same topology in our current Phylo-HMRF model. Incorporating inference of varied tree topologies (Friedman et al., 2002) will make Phylo-HMRF much more general.

To fully understand 3D genome organization evolution, it will be crucial to explore the underlying mechanisms of the different evolutionary patterns in 3D genome structure across species, e.g., in concert with the evolution of particular types of DNA sequence features that play key roles in the formation and maintenance of genome architecture and function (Sima et al., 2018; Choudhary et al., 2018; Zhang et al., 2019), which may in turn inform us about the principles of 3D genome organization. For example, we previously showed that more conserved CTCF motifs in mammalian evolution (considering motif turnover) are more likely to be involved in CTCF mediated chromatin loops (Zhang et al., 2018). Our Phylo-HMRF model has the potential to serve as a generic analytic framework to connect different evolutionary patterns of chromatin interaction with the evolution of genome sequence and function.

## Methods

### Overall framework of Phylo-HMRF for cross-species comparison of Hi-C data

We assume that a two-dimensional Hi-C contact map is given in each species, where each entry of the map represents the contact frequency between the corresponding two genomic loci. We use the human genome as the reference and align the contact pairs of genomic loci of the other species to the human genome. As a consequence, Hi-C contact maps of the other species are equivalently aligned to the human genome to be comparable. In this study, we compare the multi-species Hi-C contact frequencies in the syntenic regions genome-wide (synteny blocks were identified based on inferCARs (Ma et al., 2006)), in order to focus on the 3D genome changes that are not resulted from large-scale genome rearrangements. We then obtain a multi-species contact map **I** *∈* ℝ^*n×n×d*^, where *n* is the number of loci in the studied region on the reference genome, and *d* is the number of the species. **I** can be represented by a graph ***𝒢*** = (***𝒱***,***ℰ***), where ***𝒱*** and ***ℰ*** represent the set of nodes and the set of edges, respectively. Each node corresponds to a position in **I**, i.e., the contact between a pair of genomic loci. The number of nodes is *N* = *n × n*. We also denote ***𝒱*** as the set of indices of the nodes in ***𝒢***, i.e., ***𝒱*** = {1, …, *N*}. The *i*-th node is associated with a random variable *X*_*i*_ *∈* ℝ^*d*^ representing the multi-species observations on this node, *i ∈* ***𝒱***. The *k*-th element of *X*_*i*_ (*k* = 1, …, *d*) is the aligned contact frequency measurement of the *k*-th species between the corresponding two genomic loci. If two positions in the multi-species contact map are adjacent, there is an edge between the corresponding nodes in ***𝒢***.

Using an HMRF model, we assume that each node in ***𝒢*** is also associated with a random variable *Y*_*i*_ *∈ S* = {1, …, *M*}, representing the unknown hidden state of this node, *i ∈* ***𝒱***. *S* is the set of hidden states. For each configuration of *Y*, *X*_*i*_ follows a conditional probability distribution *p*(*x*_*i*_*|y*_*i*_), which is the emission probability distribution, and *X* = {*X*_*i*_} _*i∈****V***_ is the observable random field or emitted random field. The hidden state *Y*_*i*_ reflects different evolutionary patterns of chromatin contact frequency across species, e.g., some regions in **I** may exhibit conserved high (or low) contact frequency across species, while some may have lineage-specific high or low contact frequency. The spatial information is embedded in the MRF with the constraints on the hidden states of neighboring nodes. The neighboring nodes are expected to be more likely to have similar hidden states.

Phylo-HMRF estimates the evolutionary patterns by inferring the hidden states **y** = {*y*_*i*_}_*i∈****𝒱***_ from the observations **x** = {*x*_*i*_}_*i∈****𝒱***_, using the assumption that there are spatial dependencies between adjacent nodes in the graph ***𝒢***. In Phylo-HMRF, each hidden state *Y*_*i*_ = *l* is associated with a phylogenetic model *ψ*_*l*_. We therefore define the Phylo-HMRF model as **h** = (*S, ψ, β*), where *S* is the set of states, *ψ* is the set of phylogenetic models associated with the states, and *β* contains the pairwise potential parameters, respectively. Suppose 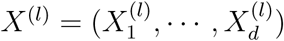 represent the values of leaf nodes of the phylogenetic tree associated with the *l*-th phylogenetic model *ψ*_*l*_, *l* = 1, …, *M*. The emission probability of each state is *p*(*X|ψ*_*l*_), which is determined by the phylogenetic model underlying this state. The joint probability of the graph ***𝒢*** is:

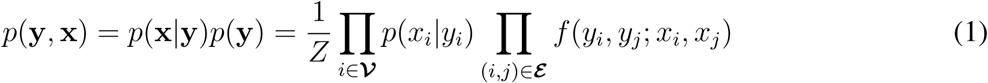

where *Z* is the normalization constant, *p*(*x*_*i*_*|y*_*i*_) is the emission probability function, which measures the probability that the local observation is generated from a certain hidden state, and *f*(·) is the compatibility function which measures the consistency of hidden states between the neighboring nodes. The joint probability can be transformed into the energy function by taking the negative logarithm of the joint probability. In this work, given the observations across species, the energy function can be defined as:

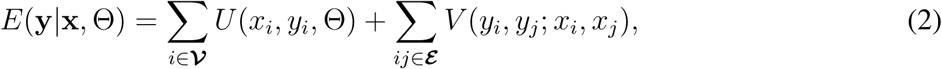

where *U* (*x*_*i*_, *y*_*i*_) is the unary potential function encoding local compatibility between observations and hidden states with model parameters Θ, and *V* (*y*_*i*_, *y*_*j*_) is the pairwise potential function encoding neighborhood information, respectively. We have that *f* (*y*_*i*_, *y*_*j*_; *x*_*i*_, *x*_*j*_) ∝ exp (*-V* (*y*_*i*_, *y*_*j*_; *x*_*i*_, *x*_*j*_)). If we take into consideration the effect of the difference between features of neighboring nodes on the pairwise potential, *V* (*y*_*i*_, *y*_*j*_; *x*_*i*_, *x*_*j*_) and the compatibility function *f* (·) will depend not only on the labels of the neighboring nodes, but also on their features or observations. We minimize the energy function to estimate the hidden states **y**. By minimizing the energy function we maximize the joint probability of the graph equivalently. As the model parameters Θ are unknown, we estimate **y** and Θ simultaneously. The objective function is:

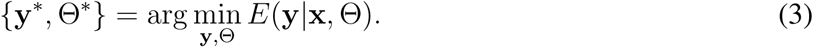

### Phylo-HMRF model with Ornstein-Uhlenbeck process

#### Ornstein-Uhlenbeck process assumptions

In Phylo-HMRF, we model the continuous traits with the Ornstein-Uhlenbeck (OU) process. The OU process is a stochastic Gaussian process that extends the Brownian motion (Felsenstein, 1985; Pagel, 1999; Freckleton, 2012) with the trend towards equilibrium around optimal values (Hansen, 1997; Butler and King, 2004; Hansen et al., 2008). The OU process has been recently used to model the evolution of genomic features (Rohlfs et al., 2013; Brawand et al., 2011; Naval-Sánchez et al., 2015; Chen et al., 2019; Yang et al., 2018). In our previous work (Yang et al., 2018), we found that the OU process has clear advantages in performance as compared to the simpler Brownian motion model. Therefore, we utilize the OU processes to realize the phylogenetic models in Phylo-HMRF.

For the observation of a lineage 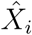, the OU process can be represented as the following (Hansen, 1997; Butler and King, 2004):

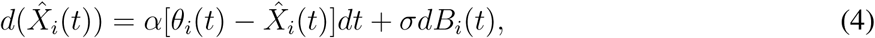

where 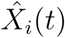 represents the observation of 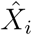 at time point *t, B*_*i*_(*t*) represents the Brownian motion, and *α, θ*_*i*_ and *σ* are parameters that represent the selection strength, the optimal value and the fluctuation intensity of Brownian motion, respectively.

For multi-species observations, based on the model assumptions of the OU process, the expectation, variance, and covariance of the observations of species can be computed given the phylogenetic tree. Suppose that *X*_*p*_ is the ancestor of species *X*_*i*_, and *X*_*a*_ is the common ancestor of species *X*_*i*_ and *X*_*j*_. We have (Butler and King, 2004; Rohlfs et al., 2013):

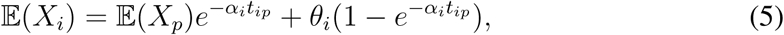

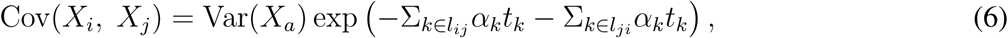

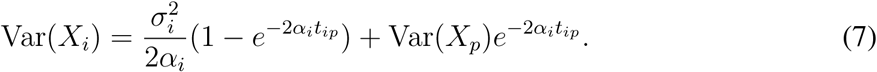

where *t*_*ip*_, *t*_*k*_ represent evolution time along corresponding branches in the phylogenetic tree, respectively. In the Phylo-HMRF model **h** = (*S, ψ, β*), *ψ*_*l*_ is defined as *ψ*_*l*_ = (*θ*_*l*_, *α*_*l*_, *σ*_*l*_, *τ*_*l*_, *b*_*l*_), 1 *≤ l ≤ M*, where *θ*_*l*_, *α*_*l*_, *σ*_*l*_ denote the optimal values, the selection strengths, and the Brownian motion intensities of the corresponding OU model, respectively, and *τ*_*l*_, *b*_*l*_ represent the topology of the phylogenetic tree, the branch lengths, respectively. *M* is the number of states. We assume that the phylogenetic tree topology is identical across different states. For the phylogenetic tree of a hidden state, we allow varied selection strengths and Brownian motion intensities along different branches and varied optimal values at the interior nodes or leaf nodes. Thus each branch is assigned a selection strength and a Brownian motion intensity, and each node is assigned an optimal value as parameters. Suppose there are *r* branches in the tree. We have 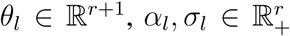, where values in *α*_*l*_ and *σ*_*l*_ are non-negative. According to the actual problem studied, *ψ*_*l*_ can be specialized to different evolutionary models. We focus on the OU processes in this work.

#### Model parameter estimation and hidden states inference

We use the Expectation-Maximization (EM) algorithm (Dempster et al., 1977; Zhang et al., 2001) for parameter estimation in our model. Zhang et al. (2001) developed the HMRF-EM algorithm where EM is adapted to estimate a HMRF model with several justified assumptions and approximations, including the pseudo-likelihood assumption (Geman and Graffigne, 1986) and mean-field approximation (Celeux et al., 2003; Zhang, 1992). The original HMRF-EM algorithm uses the multivariate Gaussian distribution as the emission probability function of a hidden state. The main difference is that in our method we use the OU processes to model the emission probability in the HMRF. Also, we utilize the Graph Cuts algorithm (Adelson and Bergen, 1991; Boykov et al., 2001) for hidden state estimation given estimates of model parameters.

Let Θ be the model parameters. Suppose Θ^*g*^ is the current estimate of model parameters. The EM algorithm aims to maximize the expectation of the complete-data log likelihood, which is defined as the *Q* function: *Q*(Θ, Θ^*g*^) = 𝔼[log *p*(**x**, **y***|*Θ)*|***x**, Θ^*g*^]. Using pseudo-likelihood approximation (Geman and Graffigne, 1986) and mean-field approximation (Celeux et al., 2003; Zhang, 1992), the *Q* function is derived as (details in Supplementary Methods):

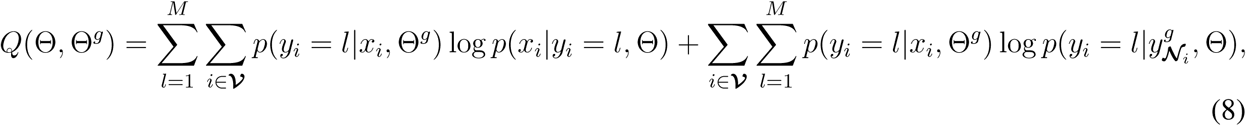

where the two parts of the *Q* function encode the unary potential and the pairwise potential of the HMRF of ***𝒢***, respectively. *p*(*y*_*i*_ = *l|x*_*i*_, Θ^*g*^) is posterior probability of each sample assigned to a hidden state given the current model parameter estimates. Using the Markov property of HMRF (Zhang et al., 2001), we have:

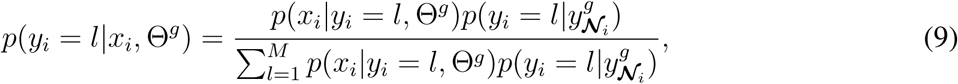

where ***𝒩***_*i*_ denotes the set of nodes that are neighbors of node *i* in ***𝒢***. We calculate *p*(*x*_*i*_*|y*_*i*_, Θ^*g*^) based on the OU process assumption (Supplementary Methods).

Let *V* (*y*_*i*_, *y*_*j*_) be the pairwise potential on a pair of adjacent nodes (*y*_*i*_, *y*_*j*_). We have:

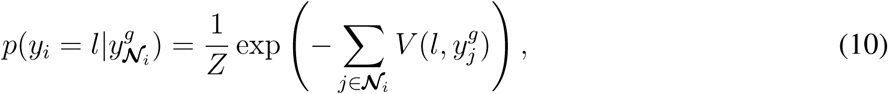

where *Z* is the normalization constant. We can adopt different definitions of the pairwise potential *V* (*y*_*i*_, *y*_*j*_) (Supplementary Methods). The definition we use takes into consideration the difference between features of the adjacent vertices in imposing the penalty on inconsistent states of the neighbors:

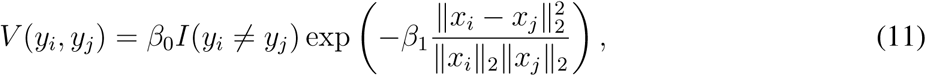

where *β*_0_, *β*_1_ are predefined adjustable regularization coefficients. Based on the definition of the pairwise potential, 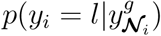 does not depend on the OU model parameters.

Let 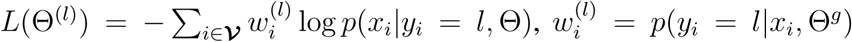. We perform parameter estimation for each of the possible states. In each Maximization-step (M-step), the objective function of a given state *l* is defined as (details in Supplementary Methods):

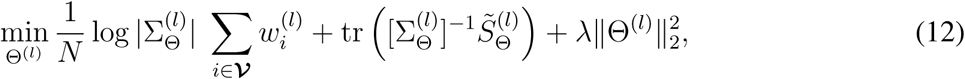

where 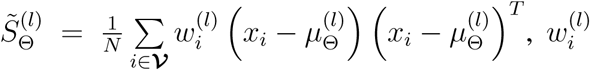 is defined as above, Θ^(*l*)^ represents the phylogenetic model parameters associated with state *l*, and *λ* is the regularization coefficient of the *l*_2_-norm regularization that is used to reduce overfitting of model (Supplementary Methods). We have that Θ^(*l*)^ = {*θ*_*l*_, *α*_*l*_, *σ*_*l*_}, where *θ*_*l*_, *α*_*l*_, *σ*_*l*_ represent the optimal values, the selection strengths, and the Brownian motion intensities of the phylogenetic tree model associated with hidden state *l*, respectively. As described previously, the OU model of state *l* is *ψ*_*l*_ = (*θ*_*l*_, *α*_*l*_, *σ*_*l*_, *τ*_*l*_, *b*_*l*_), 1 *≤ l ≤ M*. We assume that *τ*_*l*_ is given. If the branch lengths *b*_*l*_ are unknown, we perform the transformation 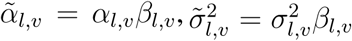 to present the combined effect of the branch length and the selection or Brownian motion parameters along this branch. Here *β*_*l,v*_ represents the length of the branch from the parent of node *v* to node *v* in the phylogenetic tree of state *l*. Then 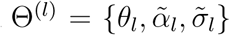, where 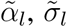 are the transformed selection strengths and the transformed Brownian motion intensities, respectively.

In Phylo-HMRF, the overall steps of the OU-model embedded HMRF-EM algorithm is as follows.

1. *Initialize the model parameter*. We perform *K*-means clustering on the samples. The clustering results are used to assign initial hidden states to the samples. For each cluster, we estimate the OU model parameters using Maximum Likelihood Estimation (MLE) (Supplementary Methods). The estimated model parameters are used as initialization of the OU model parameters for each hidden state.
2. *Estimate the hidden states given current parameter estimates*. We seek an approximate solution to the optimization problem

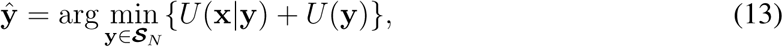

where *U* (**x***|***y**) and *U* (**y**) are the total unary potential and total pairwise potential of ***𝒢***, respectively.
3. *Calculate posterior probability distribution*. In each Expectation-step (E-step), given the current estimated model parameters Θ^*g*^ and the estimated state configuration in the previous step, we compute *p*(*y*_*i*_ = *l|x*_*i*_, Θ^*g*^) using Eq. (9, 10).
4. *Estimate model parameters by solving the optimization problem with OU models embedded*. In each M-step, we solve the MLE problem in (12) to update the parameters 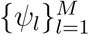.
5. *Repeat step 2-4 until convergence is reached or the maximum number of iterations is reached*.

In Phylo-HMRF, given the current estimated model parameters, we use the Graph Cuts algorithm to estimate the hidden states in Step 2 (Supplementary Methods). Graph Cuts algorithms seek to approximate solutions to an energy minimization problem by solving a max-flow/min-cut problem in a graph (Boykov et al., 2001; Boykov and Kolmogorov, 2004). Graph Cuts algorithms have been effectively used in image segmentation applications. Studies have shown that for binary image segmentation, finding a min-cut is equivalent to finding the maximum of posterior *p*(**y***|***x**) (Boykov et al., 2001). For multiple labels, the multi-labeling problem can be converted to a sequence of binary-labeling problem by *α*-expansion or *α*-*β* swap algorithms (Boykov et al., 2001). The solution is an approximate solution in the multi-labeling problem that has been shown to be a strongly probable local minima (Boykov et al., 2001).

## Acknowledgement

This work was supported in part by National Institutes of Health grant R01HG007352 (J.M.), National Institutes of Health Common Fund 4D Nucleome Program grants U54DK107965 (J.M.) and U54DK107977 (B.R.), National Institutes of Health Director’s Early Independence Award DP5OD023071 (J.D.), and National Science Foundation grant 1717205 (J.M.). The authors would like to thank members of Jian Ma’s laboratory for helpful comments to improve the manuscript.

## Author Contributions

Conceptualization, J.M.; Methodology, Y.Y., J.M.; Software, Y.Y.; Resources, B.R., J.D.; Visualization, Y.Z.; Investigation, Y.Y., Y.Z., B.R., J.D., and J.M.; Writing – Original Draft, Y.Y., and J.M.; Writing – Review & Editing, Y.Y., Y.Z., B.R., J.D., and J.M.; Funding Acquisition, B.R., J.D., J.M.

