## Supplemental Information for "Comparing 3D genome organization in multiple species using Phylo-HMRF"

#### A Supplementary Methods

##### Detailed Phylo-HMRF with OU process

In Phylo-HMRF, we embed the OU model, which is a continuous-trait phylogenetic evolutionary model into the the emission probability function of the hidden Markov random field (HMRF) model. Suppose $\Theta$ represents the model parameters, and $\Theta^g$ represents the current estimate of model parameters. The EM algorithm computes the expectation of the complete-data log likelihood, which is defined as the $Q$ function $Q(\Theta, \Theta^g)$. We have:

$$Q(\Theta, \Theta^g) = \mathbb{E}[\log p(\mathbf{x}, \mathbf{y}|\Theta)|\mathbf{x}, \Theta^g] = \sum_{\mathbf{y} \in \boldsymbol{\mathcal{S}}_N} p(\mathbf{y}|\mathbf{x}, \Theta^g) \log p(\mathbf{x}, \mathbf{y}|\Theta), \tag{14}$$

where $\mathbf{x}$ are the observations, $\mathbf{y}$ are the hidden states, and $\boldsymbol{\mathcal{S}}_N$ is the set of all the possible state configurations of size $N$. $N$ is the sample size. We have:

$$Q(\Theta, \Theta^g) = \sum_{\mathbf{y} \in \boldsymbol{\mathcal{S}}_N} p(\mathbf{y}|\mathbf{x}, \Theta^g)[\log p(\mathbf{x}|\mathbf{y}, \Theta) + \log p(\mathbf{y}|\Theta)], \tag{15}$$

where $p(\mathbf{y}|\Theta)$ represents the probability of a hidden state configuration over the whole graph $\boldsymbol{\mathcal{G}}$. Using pseudo-likelihood approximation (Geman and Graffigne, 1986), we can approximate $p(\mathbf{y}|\Theta)$ with:

$$p(\mathbf{y}|\Theta) = \prod_{i \in \boldsymbol{\mathcal{V}}} p(y_i|y_{\boldsymbol{\mathcal{N}}_i}, \Theta). \tag{16}$$

where $\boldsymbol{\mathcal{N}}_i$ denote the set of neighboring nodes of the node $i$. Then we have:

$$\begin{aligned}
Q(\Theta, \Theta^g) &= \sum_{\mathbf{y} \in \boldsymbol{\mathcal{S}}_N} p(\mathbf{y}|\mathbf{x}, \Theta^g)[\log p(\mathbf{x}|\mathbf{y}, \Theta) + \log p(\mathbf{y}|\Theta)] \\
&= \sum_{\mathbf{y} \in \boldsymbol{\mathcal{S}}_N} p(\mathbf{y}|\mathbf{x}, \Theta^g)[\sum_{i \in \boldsymbol{\mathcal{V}}} \log p(x_i|y_i, \Theta) + \sum_{i \in \boldsymbol{\mathcal{V}}} \log p(y_i|y_{\boldsymbol{\mathcal{N}}_i}, \Theta)]] \\
&= \sum_{\mathbf{y} \in \boldsymbol{\mathcal{S}}_N} \sum_{i \in \boldsymbol{\mathcal{V}}} p(\mathbf{y}|\mathbf{x}, \Theta^g) \log p(x_i|y_i, \Theta) + \sum_{\mathbf{y} \in \boldsymbol{\mathcal{S}}_N} \sum_{i \in \boldsymbol{\mathcal{V}}} p(\mathbf{y}|\mathbf{x}, \Theta^g) \log p(y_i|y_{\boldsymbol{\mathcal{N}}_i}) \\
&= \sum_{i \in \boldsymbol{\mathcal{V}}} \sum_{l=1}^{M} \sum_{q_{-i} \in \boldsymbol{\mathcal{S}}_{N-1}} p(y_i = l, y_{-i}|\mathbf{x}, \Theta^g) \log p(x_i|y_i = l, \Theta) + \sum_{\mathbf{y} \in \boldsymbol{\mathcal{S}}_N} \sum_{i \in \boldsymbol{\mathcal{V}}} p(\mathbf{y}|\mathbf{x}, \Theta^g) \log p(y_i|y_{\boldsymbol{\mathcal{N}}_i}) \\
&= \sum_{l=1}^{M} \sum_{i \in \boldsymbol{\mathcal{V}}} p(y_i = l|\mathbf{x}, \Theta^g) \log p(x_i|y_i, \Theta) + \sum_{i \in \boldsymbol{\mathcal{V}}} \sum_{\mathbf{y} \in \boldsymbol{\mathcal{S}}_N} p(\mathbf{y}|\mathbf{x}, \Theta^g) \log p(y_i|y_{\boldsymbol{\mathcal{N}}_i}).
\end{aligned} \tag{17}$$

Let $x_{\boldsymbol{\mathcal{N}}_i}$ denote the neighbors of $x_i$, and $y_{\boldsymbol{\mathcal{N}}_i}$ denote the state configuration of $x_{\boldsymbol{\mathcal{N}}_i}$.

Calculating $\sum_{i \in \boldsymbol{\mathcal{V}}} \sum_{\mathbf{y} \in \boldsymbol{\mathcal{S}}_N} p(\mathbf{y}|\mathbf{x}, \Theta^g) \log p(y_i|y_{\boldsymbol{\mathcal{N}}_i})$ requires computing $p(\mathbf{y}|\mathbf{x}, \Theta^g)$ and $\log p(y_i|y_{\boldsymbol{\mathcal{N}}_i})$ over all the possible configurations $\mathbf{y} \in \boldsymbol{\mathcal{S}}_N$, which is computationally intractable. By mean-field approximation (Celeux et al., 2003; Zhang, 1992), we can use estimated hidden states $y^g_{\boldsymbol{\mathcal{N}}_i}$ from the previous iteration of the HMRF-EM algorithm to approximate $p(y_i|y_{\boldsymbol{\mathcal{N}}_i})$, which can simplify the computation of the $Q$ function. Then we have:

$$\sum_{i\in\boldsymbol{\mathcal{V}}}\sum_{\mathbf{y}\in\boldsymbol{\mathcal{S}}_N} p(\mathbf{y}|\mathbf{x},\Theta^g)\log p(y_i|y^g_{\boldsymbol{\mathcal{N}}_i}) = \sum_{i\in\boldsymbol{\mathcal{V}}}\sum_{l=1}^{M}\sum_{q_{-i}\in\boldsymbol{\mathcal{S}}_{N-1}} p(y_i=l,y_{-i}|\mathbf{x},\Theta^g)\log p(y_i=l|y^g_{\boldsymbol{\mathcal{N}}_i})$$
$$= \sum_{l=1}^{M}\sum_{i\in\boldsymbol{\mathcal{V}}} p(y_i=l|x_i,\Theta^g)\log p(y_i=l|y^g_{\boldsymbol{\mathcal{N}}_i},\Theta). \tag{18}$$

Therefore,

$$Q(\Theta,\Theta^g) = \sum_{l=1}^{M}\sum_{i\in\boldsymbol{\mathcal{V}}} p(y_i=l|x_i,\Theta^g)\log p(x_i|y_i=l,\Theta) + \sum_{i\in\boldsymbol{\mathcal{V}}}\sum_{l=1}^{M} p(y_i=l|x_i,\Theta^g)\log p(y_i=l|y^g_{\boldsymbol{\mathcal{N}}_i},\Theta). \tag{19}$$

The parameters of the OU model are embedded in the term $p(x_i|y_i)$ of $Q(\Theta,\Theta^g)$. The second part of the $Q$ function encodes the pairwise potentials and does not include the OU model parameters. Therefore, the first and second parts of the $Q$ can be estimated separately.

Using the Markov property of the HMRF model (Zhang et al., 2001), we have:

$$p(y_i=l|x_i,\Theta^g) = \frac{p(x_i|y_i=l,\Theta^g)p(y_i=l|y^g_{\boldsymbol{\mathcal{N}}_i})}{\sum_{l=1}^{M} p(x_i|y_i=l,\Theta^g)p(y_i=l|y^g_{\boldsymbol{\mathcal{N}}_i})}. \tag{20}$$

Based on the model of the OU process, we assume the observations of observed species (leaf nodes in the phylogenetic tree) follow multivariate Gaussian distribution. We have:

$$p(x_i|y_i=l) = p(x_i|\Theta^{(l)})$$
$$= \frac{1}{(2\pi)^{d/2}|\Sigma^{(l)}_\Theta|^{1/2}}\exp\left\{-\frac{1}{2}\left(x_i-\mu^{(l)}_\Theta\right)^T[\Sigma^{(l)}_\Theta]^{-1}\left(x_i-\mu^{(l)}_\Theta\right)\right\}, \tag{21}$$
$$\log p(x|\Theta^{(l)}) \propto -\frac{1}{2}\log|\Sigma^{(l)}_\Theta| - \frac{1}{2}\left(x_i-\mu^{(l)}_\Theta\right)^T[\Sigma^{(l)}_\Theta]^{-1}\left(x_i-\mu^{(l)}_\Theta\right), \tag{22}$$

where $\Theta^{(l)}$ represent the OU model parameters associated with the $l$-th state and $d$ is the number of the observed species. The underlying phylogenetic model $\psi_l$ is embedded into $\Sigma^{(l)}_\Theta$ and $\mu^{(l)}_\Theta$ according to Eq. (5)-(7). Let $w^{(l)}_i = p(y_i=l|x_i,\Theta^g)$. We have:

$$Q(\Theta,\Theta^g) = \sum_{l=1}^{M}\sum_{i\in\boldsymbol{\mathcal{V}}} w^{(l)}_i\log p(x_i|y_i=l,\Theta) + \sum_{l=1}^{M}\sum_{i\in\boldsymbol{\mathcal{V}}} w^{(l)}_i\log p\left(y_i=l|y^g_{\boldsymbol{\mathcal{N}}_i}\right) \tag{23}$$
$$= \sum_{l=1}^{M}\sum_{i\in\boldsymbol{\mathcal{V}}} w^{(l)}_i\left[-\frac{1}{2}\log|\Sigma^{(l)}_\Theta| - \frac{1}{2}\left(x_i-\mu^{(l)}_\Theta\right)^T[\Sigma^{(l)}_\Theta]^{-1}\left(x_i-\mu^{(l)}_\Theta\right) + \log p\left(y_i=l|y^g_{\boldsymbol{\mathcal{N}}_i}\right)\right] + C, \tag{24}$$

where $C$ is a constant.

We perform parameter estimation for each of the possible states.

Let $L(\Theta^{(l)}) = -\sum_{i\in\boldsymbol{\mathcal{V}}} w^{(l)}_i\log p(x_i|y_i=l,\Theta)$. We have:

$$L(\Theta^{(l)}) = \frac{1}{2}\log|\Sigma^{(l)}_\Theta|\sum_{i\in\boldsymbol{\mathcal{V}}} w^{(l)}_i + \frac{1}{2}\sum_{i\in\boldsymbol{\mathcal{V}}}\left(x_i-\mu^{(l)}_\Theta\right)^T[\Sigma^{(l)}_\Theta]^{-1}\left(x_i-\mu^{(l)}_\Theta\right)w^{(l)}_i. \tag{25}$$

Therefore, the first part of the negative $Q$ function with respect to a given state $l$ can be represented as:

$$\tilde{L}(\Theta^{(l)}) = \frac{1}{N} \log |\Sigma_{\Theta}^{(l)}| \sum_{i \in \boldsymbol{\mathcal{V}}} w_i^{(l)} + \mathrm{tr}\left( [\Sigma_{\Theta}^{(l)}]^{-1} \tilde{S}_{\Theta}^{(l)} \right), \tag{26}$$

where $\tilde{S}_{\Theta}^{(l)} = \frac{1}{N} \sum_{i \in \boldsymbol{\mathcal{V}}} w_i^{(l)} \left( x_i - \mu_{\Theta}^{(l)} \right) \left( x_i - \mu_{\Theta}^{(l)} \right)^T$, and $\Theta^{(l)}$ represents the phylogenetic model parameters associated with state $l$. We assume the phylogenetic tree topology $\tau_l$ is given. The branch lengths $b_l$ can be combined in effect to $\alpha_l$ and $\sigma_l$. As we allow varied selection strength and Brownian motion intensity of the OU model along each branch of the phylogenetic tree, and allow varied optimal values on each tree node, there are many OU model parameters to estimate, which may result in overfitting of the model if the sample size is not large enough. We apply $\ell_2$-norm regularization to the parameters $\Theta^{(l)}$ to reduce model overfitting. In each M-step, the objective function of a given state $l$ is defined as:

$$\min_{\Theta^{(l)}} \frac{1}{N} \log |\Sigma_{\Theta}^{(l)}| \sum_{i \in \boldsymbol{\mathcal{V}}} w_i^{(l)} + \mathrm{tr}\left( [\Sigma_{\Theta}^{(l)}]^{-1} \tilde{S}_{\Theta}^{(l)} \right) + \lambda \|\Theta^{(l)}\|_2^2, \tag{27}$$

where $\tilde{S}_{\Theta}^{(l)} = \frac{1}{N} \sum_{i \in \boldsymbol{\mathcal{V}}} w_i^{(l)} \left( x_i - \mu_{\Theta}^{(l)} \right) \left( x_i - \mu_{\Theta}^{(l)} \right)^T$, $w_i^{(l)}$ is defined as above, $\Theta^{(l)}$ represents the phylogenetic model parameters associated with state $l$. $\lambda$ is the regularization coefficient. We define $\lambda = \lambda_0 / \sqrt{N}$. $\lambda_0$ can be predefined. We choose $\lambda_0 = 4.0$ in both the simulation study and real data study. The same regularization coefficient was adopted in (Yang et al., 2018) and the value of $\lambda_0 = 4.0$ was used. We found that the performance of the model was not sensitive to the choice of $\lambda_0$ within a range. We therefore use the same choice of $\lambda_0$ in Phylo-HMRF.

For the second part of the $Q$ function, we can have different definitions of the pairwise potential $V(y_i, y_j)$. We will consider two definitions. One definition is:

$$V(y_i, y_j) = \beta_0 I(y_i \neq y_j), \tag{28}$$

where $\beta_0$ is a predefined adjustable regularization coefficient, which can also be considered as pairwise potential parameter.

The second definition takes into consideration the difference of features of the adjacent vertices in imposing the penalty on inconsistent states of the neighbors.

$$V(y_i, y_j) = \beta_0 I(y_i \neq y_j) \exp\left( -\beta_1 \frac{\|x_i - x_j\|_2^2}{\|x_i\|_2 \|x_j\|_2} \right), \tag{29}$$

where $\beta_0$ and $\beta_1$ are predefined adjustable regularization coefficients, which can also be considered as pairwise potential parameters. In Phylo-HMRF we mainly use the second definition. $\beta_0$ and $\beta_1$ can either be estimated as model parameters or predefined. In many applications pairwise potential parameters are usually estimated through a number of trials and predefined. The pairwise potential parameters can be chosen such that the pairwise potential is at the same scale of the unary potential. We choose $\beta_0 \in [1, 3]$ and $\beta_1 \in [0.1, 0.5]$ in the simulation evaluation and the real data application based on empirical observations from a simulation dataset.

#### Model initialization in Phylo-HMRF

In the OU-model embedded HMRF-EM algorithm, we need to initialize the model parameters. We follow the similar approaches in Yang et al. (2018) for parameter initialization in the EM algorithm.

For the first approach, we perform $K$-means clustering on the samples. We assign a hidden state to the samples in the same cluster. For each cluster, we estimate the OU model parameters by maximum likelihood estimation (MLE). The objective function of the MLE problem is similar to that defined in Eq. (27). The difference is that we set $w_i^{(l)} = 1$, and change $i \in \boldsymbol{\mathcal{V}}$ to the constraint $i \in \boldsymbol{\mathcal{C}}_l$, where $\boldsymbol{\mathcal{C}}_l$ represents the set of the nodes that are assigned to state $l$ by the $K$-means clustering result. The estimated model parameters are used as initialization of the OU model parameters for each hidden state. The second approach is to randomly sample the parameter values from predefined uniform distributions.

For the third approach, we use a linear combination of parameters obtained from the first and second approaches for parameter initialization. The initial parameter values are chosen as $\Theta_0 = w_1\Theta_1 + (1 - w_1)\Theta_2$, where $w_1 \in [0, 1]$, $\Theta_1$ and $\Theta_2$ are parameter estimates from the first and second approaches, respectively. In practice, we used the third approach.

#### HMRF-EM algoirthm and Graph Cuts algorithm used in Phylo-HMRF

In Phylo-HMRF, given the current estimated model parameters, we use the Graph Cuts algorithm to estimate the hidden states in Step 2 of the HMRF-EM algorithm. In step 2, we seek approximate solution to the energy minimization problem:

$$\hat{\mathbf{y}} = \arg\min_{\mathbf{y}\in\boldsymbol{\mathcal{S}}_N} \{U(\mathbf{x}|\mathbf{y}) + U(\mathbf{y})\}, \tag{30}$$

where $U(\mathbf{x}|\mathbf{y}$ and $U(\mathbf{y})$ are the unary potential and the pairwise potential, respectively. We have also shown that minimizing the energy is equivalent to maximizing the joint probability.

We seek the solution $\{\mathbf{y}^*, \Theta^*\} = \arg\max_{\mathbf{y},\Theta} E(\mathbf{y}|\mathbf{x}, \Theta)$ by alternatively performing

$$\mathbf{y}^* = \arg\min_{\mathbf{y}} E(\mathbf{y}|\mathbf{x}, \Theta^g), \tag{31}$$

and

$$\Theta^* = \arg\max_{\Theta} \mathbb{E}\left[\log p(\mathbf{x}, \mathbf{y}|\Theta)|\mathbf{x}, \Theta^g\right], \tag{32}$$

where $\Theta^g$ is the current estimates of the model parameters. We use $\mathbf{y}^*$ in computing $\mathbb{E}[\log p(\mathbf{x}, \mathbf{y}|\Theta)|\mathbf{x}, \Theta^g]$ with the mean-field approximation.

We use Graph Cuts algorithm for the first stage (Eq. (31)) and use EM algorithm for the second stage (Eq. (32)). The energy minimization problem for MRF (Eq. (31)) is known to be NP-hard. Graph Cuts algorithms can effectively seek approximate solutions to an energy minimization problem by solving a max-flow/min-cut problem in a graph (Boykov et al., 2001). We define the unary cost and the pairwise cost of the graph $\boldsymbol{\mathcal{G}}$ to utilize the Graph Cuts algorithm. The unary cost corresponds to the unary potential, which is:

$$U(x_i|y_i, \Theta^g) \propto -\log(p(x_i|y_i, \Theta^g). \tag{33}$$

The pairwise cost corresponds to the pairwise potentials. We compute the edge weights in $\boldsymbol{\mathcal{G}}$ by calculating:

$$\bar{w}_{ij} = \exp\left(-\beta_1 \frac{\|x_i - x_j\|_2^2}{\|x_i\|_2\|x_j\|_2}\right), \tag{34}$$

where $\beta_1$ is an coefficient, and provide a pairwise state transition cost matrix $\bar{V} \in \mathbb{R}^{M\times M}$, where $M$ is the number of hidden states and $\bar{V}_{ij}$ represents the penalty on $y_j \neq y_i$ for an directed edge $(i \leftarrow j)$. In our problem, $\boldsymbol{\mathcal{G}}$ is an undirected graph, and we simplify $\bar{V}$ as $\bar{V}_{ij} = \beta_0$, $i, j = 1, \cdots, M$. However, in more complicated problem settings, we can realize $\bar{V}$ with varied elements $\bar{V}_{ij}$ and estimate the elements as model parameters. We use the GCO library to perform the Graph Cuts algorithm (Boykov and Kolmogorov, 2004; Boykov et al., 2001; Kolmogorov and Zabih, 2004).

#### Approach to generating the simulated datasets

In the simulation evaluation, we suppose that the samples in simulated dataset correspond to nodes in a graph. Similar to a Hi-C contact map, $\boldsymbol{\mathcal{G}}$ has 2D lattice structure of size $n \times n$, where each vertex is associated with a sample. The samples thus represent features of vertices in $\boldsymbol{\mathcal{G}}$. Each node in $\boldsymbol{\mathcal{G}}$ can be assigned 2D coordinates based on its position in the graph. Let $\boldsymbol{\mathcal{N}}_i$ denote the set of neighboring nodes of the node $i$, i.e., the nodes that are connected to node $i$ in $\boldsymbol{\mathcal{G}}$. We use 8-connected neighborhood system. Suppose the node $i$ has coordinates $(c_{i_1}, c_{i_2})$. Then the nodes with coordinates $(c_{i_1}, c_{i_2} \pm 1)$, $(c_{i_1} \pm 1, c_{i_2})$, $(c_{i_1} - 1, c_{i_2} \pm 1)$, and $(c_{i_1} + 1, c_{i_2} \pm 1)$ are the neighbors of node $i$.

In simulation study I, for each simulated dataset, we first simulate a configuration of the hidden states of the samples by simulating an MRF through Gibbs sampling (Geman and Geman, 1984). We assume that each sample has a hidden state. Each hidden state is associated with an emission probability function. The hidden states of the samples are assumed to be from an MRF. We use the Markov property:

$$p(y_i|y_{-i}) = p(y_i|y_{\boldsymbol{\mathcal{N}}_i}), \tag{35}$$

where $y_{-i}$ represents the hidden states of all the nodes other than the $i$th node, i.e., $y_{-i} = \{y_j, j \in \boldsymbol{\mathcal{V}}, j \neq i\}$. $y_{\boldsymbol{\mathcal{N}}_i}$ represents the hidden states of the neighbors of node $i$.

We randomly initialize the hidden state configuration of the $N = n \times n$ samples at time step $t = 0$. The hidden state $y_i^{(t)}$ of sample $i$ at time step $(t+1)$ is sampled from the probabilistic distribution $p(y_i|y_{\boldsymbol{\mathcal{N}}_i}^{(t-1)})$, $y_i \in S = \{1, \cdots, M\}$. $p(y_i = l|y_{\boldsymbol{\mathcal{N}}_i}^{(t-1)})$ is calculated using Eq. (10), $l \in S$. We use the first definition of pairwise potential and use $\beta_0 = 2$. We repeat this sampling process until the maximum of time steps to take $T$ is reached. We use $\mathbf{y}^{(T)} = \{y_i^{(T)}\}_{i \in \boldsymbol{\mathcal{V}}}$ as the hidden states of the samples. We then simulate observations of the samples based on the hidden states using the emission probability functions $p(x_i|y_i, \theta_{y_i})$, where $\theta_{y_i}$ represents parameters of the emission probability distribution of hidden state $y_i$. We assume the emission probability function of each hidden state is a multivariate Gaussian distribution. Suppose the observations are $x_i \in \mathbb{R}^d$, $i \in \boldsymbol{\mathcal{V}}$. Let $d$ be the number of species. $x_i$ then represents the multi-species observations. We assume that each of the Gaussian distribution is associated with a different OU model. For each hidden state, we randomly sample the OU model parameters selection strength and Brownian motion on each branch from uniform distribution Unif$[0, 1]$, and sample the optimal values from normal distribution $\boldsymbol{\mathcal{N}}(0.5, 0.25)$. We then use OU model parameters to calculate the Gaussian distribution parameters $\theta_l$, $l \in S$, and simulate samples based on the hidden states and the corresponding multi-variate Gaussian distributions $p(x_i|y_i, \theta_{y_i})$. We use $n = 500$, $N = 250000$, $d = 4$, $M = 10$, and use the same topology of the phylogenetic tree as we use in real data analysis for the OU models that are associated with the Gaussian distributions. In simulation study II, we use the same eight set of hidden states simulated in simulation study I, but simulate OU model parameters with different parameter settings. We randomly sample the OU model parameters selection strength and Brownian motion intensity on each branch from uniform distribution Unif$[0, 1.5]$ and sample the optimal values from normal distribution $\boldsymbol{\mathcal{N}}(1, 0.5)$. We also use $n = 500$, $N = 250000$, $d = 4$, $M = 10$.

We assume that the data simulation process is hidden from us. When we applied Phylo-HMRF to the simulated datasets, we used the second definition of the pairwise potential by considering the feature difference of adjacent nodes, and used $\beta_0 = 1$, $\beta_1 = 0.1$. Therefore, the parameter settings we used to implement HMRF-EM are different from the simulation parameter settings, which can better test whether the model has robust capability. We also tried varied parameters $\beta_0 = 1.5$, and $\beta_0 = 2$ and tested the performance of Phylo-HMRF. We found that Phylo-HMRF still maintains higher accuracy than the other methods and the performance is improved to a moderate level, demonstrating the robustness of Phylo-HMRF. We only report the results obtained with $\beta_0 = 1$ in the performance evaluation.

#### Other methods compared in the simulation evaluation

In the simulation evaluation, we compared Phylo-HMRF with the Gaussian-HMRF method (Zhang et al., 2001), the Gaussian Mixture Model (GMM), the *K*-means clustering method, and two image segmentation methods SLIC and Quick Shift in state estimation. To utilize the image segmentation methods, we consider the combined multi-species Hi-C contact map as an image, and consider the features of each species as one color channel of the image. We normalize the features of each species to be in the range [0,1] accordingly, which is the scale of a color channel, to prepare the input for SLIC and Quick Shift. For the segmentation results, we consider segments with the same label as the same state. SLIC performs *K*-means clustering in the joint space of color information and spatial coordinates over an image. Quick Shift is approximation of the Mean Shift algorithm (Comaniciu and Meer, 2002) with kernel methods utilized, performing mode seeking in segmenting an image. We use the scikit-image package (Van der Walt et al., 2014) that includes implementation of the SLIC and Quick Shift algorithms. For the methods Gaussian-HMRF, GMM, and *K*-means clustering, we set the number of states to be 10, respectively, which is the number of ground truth states. For the two image segmentation methods implemented by scikit-image, there are no input arguments to set the exact number of output segments. We adjust the parameter configurations of each of the two methods such that the number of output segments is approximately 10 and comparable to the state estimation results of the other methods. For SLIC, we use an argument to set the approximate number of segments and adjust the other parameters to have the number of output segments approximating 10. For Quick Shift, there is no argument to set an exact or approximate number of output segments. We then tune the input parameters to have the number of output segments approximating 10.

#### Performance metrics in the simulation evaluation

We evaluated the accuracy of Phylo-HMRF in estimating hidden states by comparing the predicted states with the ground truth states in the simulation evaluation, using evaluation metrics Normalized Mutual Information (NMI), Adjusted Mutual Information (AMI), Adjusted Rand Index (ARI), Precision, Recall, and $F_1$ score (Manning et al., 2008; Vinh et al., 2010). These metrics compare two partitions of a set. Suppose $X = \{x_1, \cdots, x_N\}$ are the samples. Suppose $\Omega = \{\omega_1, \cdots, \omega_K\}$ and $C = \{c_1, \cdots, c_M\}$ are the predicted partition of the samples and the ground truth partition of the samples, respectively. The mutual information (MI) between $\Omega$ and $C$ is $I(\Omega; C) = \sum_{k=1}^{K} \sum_{j=1}^{M} P(\omega_k, c_j) \log \frac{P(\omega_k, c_j)}{P(\omega_k)P(c_j)}$. The NMI between $\Omega$ and $C$ is:

$$NMI(\Omega; C) = \frac{I(\Omega; C)}{[H(\Omega) + H(C)]/2} \tag{36}$$

where $H(\Omega)$ and $H(C)$ are the entropies of $\Omega$ and $C$, respectively. $H(\Omega) = -\sum_{k=1}^{K} P(\omega_k) \log P(\omega_k)$, $H(C) = -\sum_{j=1}^{M} P(c_j) \log P(c_j)$.

AMI corrects MI by removing the effect of agreement between two partitions that is due to chance. AMI is defined as:

$$AMI(\Omega; C) = \frac{I(\Omega; C) - \mathbb{E}[I(\Omega; C)]}{\max\{H(\Omega), H(C)\} - \mathbb{E}[I(\Omega; C)]}, \tag{37}$$

where $\mathbb{E}(I(\Omega; C))$ represents the expectation of $I(\Omega; C)$, which can be estimated based on $\Omega$ and $C$ (Vinh et al., 2010).

The Rand Index (RI) (Manning et al., 2008) also compares two partitions, which is defined as:

$$RI = \frac{TP + TN}{TP + FP + FN + TN}, \tag{38}$$

where TP (true positive), FP (false positive), FN (false negative), and TN (true negative) represent the number of sample pairs that are in the same subset in $\Omega$ and also in the same subset in $C$, the number of sample pairs that are in the same subset in $\Omega$ but in different subsets in $C$, the number of sample pairs that are in different subsets in $\Omega$ but in the same subset in $C$, and the number of sample pairs that are in different subsets in $\Omega$ and also in different subsets in $C$, respectively.

ARI corrects RI by removing the effect of agreement between partitions that is due to chance.

$$ARI = \frac{RI - \mathbb{E}[RI]}{\max\{RI\} - \mathbb{E}[RI]}, \tag{39}$$

where $\mathbb{E}(RI)$ represents the expectation of $RI$.

Precision, Recall, and $F_1$ score are defined as

$$Precision = \frac{TP}{TP + FP}, \tag{40}$$

$$Recall = \frac{TP}{TP + FN}, \tag{41}$$

$$F_1 = \frac{2Precision \times Recall}{Precision + Recall}. \tag{42}$$

#### Cross-species Hi-C data processing

We used the Hi-C data from the lymphoblastoid cells in human (GM12878) from the 4DN data portal and generated Hi-C data in lymphoblastoid cells in chimpanzee, bonobo, and gorilla. The genome assemblies used for the four species are hg38, panTro5, panPan2, and gorGor4, respectively. The genome assemblies were downloaded from the UCSC genome browser. We used the software Juicer Tools (Durand et al., 2016) to process the Hi-C sequencing reads of each of the three species to obtain the Hi-C contact pairs based on the corresponding genome assembly. Each Hi-C contact pair is a pair of reads that are mapped to the two genomic loci based on the corresponding genome assembly, representing chromatin contact between these two genomic loci. For the human GM12878 data, the Hi-C contacts file resulted from merging and processing all replicates at the 4DN data portal has much higher coverage than the data for other primate species, affecting the comparability of the Hi-C contact maps between human and the other species. We therefore performed random sampling to obtain approximately $2.9 \times 10^8$ contact pairs in human, comparable to the other species. We obtained approximately $2.9 \times 10^8$, $2.7 \times 10^8$, $2.4 \times 10^8$, and $2.9 \times 10^8$ contact pairs for human, chimpanzee, bonobo, and gorilla, respectively.

Next, we aligned the Hi-C contact pairs of the non-human species to the human genome. We mapped the aligned loci of the two ends of a contact pair in the Hi-C data of the non-human species from the original genome assembly to the human genome with reciprocal mapping using the tool liftOver (Hinrichs et al., 2006). With this conversion, the Hi-C contact maps of the four species that are computed from the aligned Hi-C contact pairs are all based on the human genome coordinates and therefore can be directly compared in the synteny blocks across species. The synteny blocks are the genome regions where the order of genome loci are preserved and there are no chromosome rearrangements greater than a certain resolution (50 kb in this study).

Next, we used Juicer Tools to generate the Hi-C contact maps of each species at the resolution of 50Kb from the .hic files of the corresponding species, performing normalization by Knight-Ruiz matrix balancing (Durand et al., 2016). For the Hi-C contact map of each species in each synteny block, we perform two-step filtering to interpolate the missing values and smooth the signals. In the first step we use the median filter for interpolation of possible missing values. For each node without signal value in the Hi-C contact map, we use the median of Hi-C contact frequencies of the 8-connected neighbors as the value assigned to the node. Median filter has the characteristic to preserve edges in image. In the second step, we apply an anisotropic diffusion filter, which is an edge-preserving filter to the whole Hi-C contact map in this synteny block to smooth the signals while maintaining the edge features, which correspond to more rapid change of Hi-C contact frequencies. After preprocessing the Hi-C map of each species, we then align the Hi-C contact maps of the four species in each synteny block to obtain a combined multi-species Hi-C contact map, where each node in the map corresponds to Hi-C contact frequencies between the corresponding pair of genome loci in the four species. As the scales of Hi-C contact frequencies in different species are different, we normalize the Hi-C contact frequencies of each species to the same scale over all the autosomes. We then perform the $\hat{x} = \log(1 + x)$ transformation to the normalized Hi-C contact signals of each species.

We focus on the comparison of Hi-C data in synteny blocks across species in this study. Therefore, we use the Hi-C contact signals within the synteny blocks as input to Phylo-HMRF, which are the subgraphs of the combined multi-species Hi-C contact map of each chromosome. The multi-species Hi-C contact map of each synteny block is symmetric. Therefore, we only keep the up-triangular part of the Hi-C contact map, and consider each entry as a sample. Each sample is a multi-dimensional feature vector, where each dimension represents the Hi-C contact frequency of the corresponding species between the corresponding pair of genomic loci specified by the coordinates of the entry in the Hi-C contact map. Hence, there are $N(N-1)/2$ samples for a synteny block of size $N$. We originally identified 90 synteny blocks in 50kb resolution on the autosomes based on inferCARs (Ma et al., 2006). There are two large size synteny blocks on chromosome 3 and chromosome 6, which exceeds 150Mb and 190Mb each. We then divided the two large synteny blocks into two parts each according to the two chromosome arms, respectively. For each divided synteny block, we still consider the interactions between the two subregions. Overall, we have 92 synteny blocks identified in the autosomes with 30,154,205 samples.

#### Initial estimation of the number of states for Phylo-HMRF

We estimated the possible number of hidden states using the *K*-means clustering before applying Phylo-HMRF to the cross-species Hi-C data. We performed *K*-means clustering to the cross-species Hi-C data with the cluster number *K* increasing from 2 to 100. We computed the Sum of Squared Error (SSE) of each clustering result, and observed how SSE changed with respect to the different choices of *K* by plotting the SSE-*K* curve (Fig. S3). We found that approximately the decreasing rate of SSE with respect to the increasing *K* slows down in the range of 15-30. Small fluctuation of the number of states around 30 dose not result in significant reduction of SSE with respect to the increase of *K*. We therefore set the number of hidden states to be 30.

#### State estimation on the Hi-C data by Phylo-HMRF

The difference of Hi-C contact frequency across species can be either resulted from genome rearrangements or other types of genome evolution. In this work, we specifically focus on changes within synteny blocks. For all the autosomes in the human genome, we run Phylo-HMRF jointly on the multiple synteny blocks of the chromosomes and identified possible different evolutionary patterns of the Hi-C contact frequencies across species in a genome-wide manner. When applying Phylo-HMRF to the Hi-C data, we use the second definition of the pairwise potential by considering the feature difference of adjacent nodes, and use $\beta_0 = 3$, $\beta_1 = 0.1$. We tested different choices of $\beta_0$, compared the hidden state estimation results to the Hi-C contact maps by observation, and chose $\beta_0 = 3$ as an estimate of $\beta_0$.

We further categorize the 30 estimated hidden states into 13 groups, as described in the Results section. Base on the Hi-C contact frequency distributions in the four species in each estimated state, we identify the states with distinctively higher or lower Hi-C contact frequency values than other states in all the four species as the C-high and C-low states, respectively. For the other states, the states showing similar feature distributions of Hi-C contact frequency in four species are annotated as C-mid and WC. The rest states are annotated as non-conserved (NC) states and we further identify the lineage-specific states where one species showing distinctive divergence in feature distribution from the other species.

After applying Phylo-HMRF to the cross-species Hi-C data for hidden state estimation results, we obtained segmentation of the cross-species Hi-C contact map, where each node is assigned a label which represents the estimated hidden state. Neighboring nodes with the same hidden state form a local segment. The segmentation results can be visualized as a color image. We then perform simple post-processing of the segmentation results, in order to obtain more smoothed segmentation which could facilitate downstream analysis.

For the segmentation resulted from state estimation in each synteny block of a chromosome, we consider it as an image and first find all the connected components in this image. A connected component in a graph is a subgraph in which every two nodes are connected by a path and each node within is not connected to external nodes outside the subgraph. Each connected component can be considered as a segment of the cross-species Hi-C contact map. Nodes in a connected component have the same estimated states. If the size of the connected component (i.e., the number of nodes in the segment) is smaller than a threshold, we query the states of all the external nodes in local neighborhood to any node in the component and use the most frequent observed state of the external neighbors to reassign states for this component. We set the threshold to be 10, and use window size of 5 to define local neighborhood surrounding a node. We performed this post-processing step once and obtained slightly smoothed segmentation results. The threshold can be adjusted and the procedure can be iterated to obtain more smoothed segmentation.

##### Alignment between boundaries of identified local-contact block patterns and TADs

For segments of estimated Hi-C evolutionary patterns on the diagonal of the Hi-C map of a synteny block, we use windows that could match the segments to identify the local-contact block patterns. Specifically, we use a sliding window with changeable size to find possible matches to the segments on the diagonal. With a window of the lower bound size located at a starting position along the diagonal, we first find the dominant estimated state within this window. If there is no dominant state, we move the window to the next position by one bin. The dominant state is defined as the state with the percentage exceeding a threshold and with the highest percentage within the window. We then increase the window size from the lower bound gradually until the percentage of the dominant state within the window decreases or the dominant state changes or disappears. If a window has the highest percentage of the dominant state and the percentage reaches the threshold (we use 0.95), we identify it as a local-contact block. We then reset the window to the lower bound size and move it to the next position by a stride that is half of the previously identified block size. We repeat the steps above until we scan all the estimated states on the diagonal of the Hi-C contact map.

To estimate the empirical distribution of the distance between random block boundaries on the diagonal and TAD boundaries, for each synteny block, we shuffle the identified local-contact block patterns on the diagonal of the cross-species Hi-C contact map 1000 times by randomly relocating them within this synteny block. For each boundary of each randomly relocated diagonal block in a shuffle, we calculate the distance between the block boundary and the nearest TAD boundary of a specific type of TAD (i.e., Arrowhead TAD or DI TAD). For each shuffle, we then compute the percentages of the diagonal blocks the distance of which to the corresponding nearest TAD boundaries fall in five distance intervals, which are 0-50Kb, 50-100Kb, 100-150Kb, 150-200Kb, >200Kb. We merge the percentages from each shuffle of diagonal block patterns as an empirical distribution for each distance interval.

### B Supplementary Figures and Tables

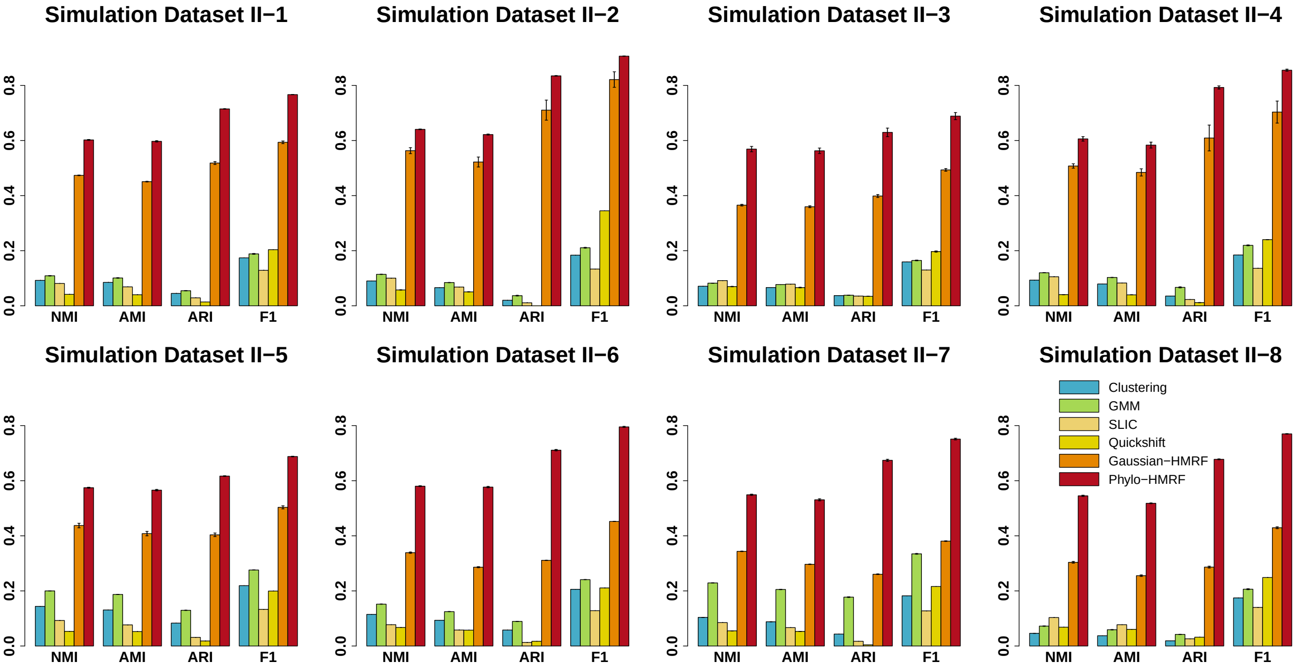


**Figure S1:** Performance evaluation of $K$-means Clustering, GMM, SLIC, Quickshift, Gaussian-HMRF, and Phylo-HMRF on eight simulation datasets in simulation study II with respect to NMI (Normalized Mutual Information), AMI (Adjusted Mutual Information), ARI (Adjusted Rand Index), and $F_1$ score. The standard error of the results from 10 runs of each method is shown as the error bar, respectively.

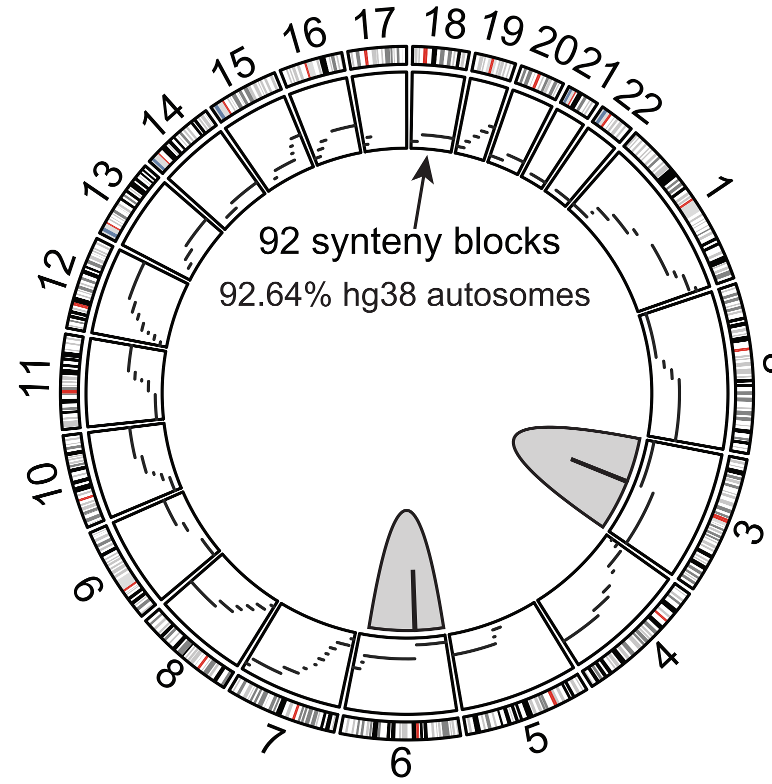


**Figure S2:** Distribution of the identified synteny blocks on the 22 autosomes of the human genome. For chromosome 3 and chromosome 6, we divide the large size synteny blocks into two parts each according to the chromosome arms. The synteny blocks cover 92.64% of human chromosomes 1-22.

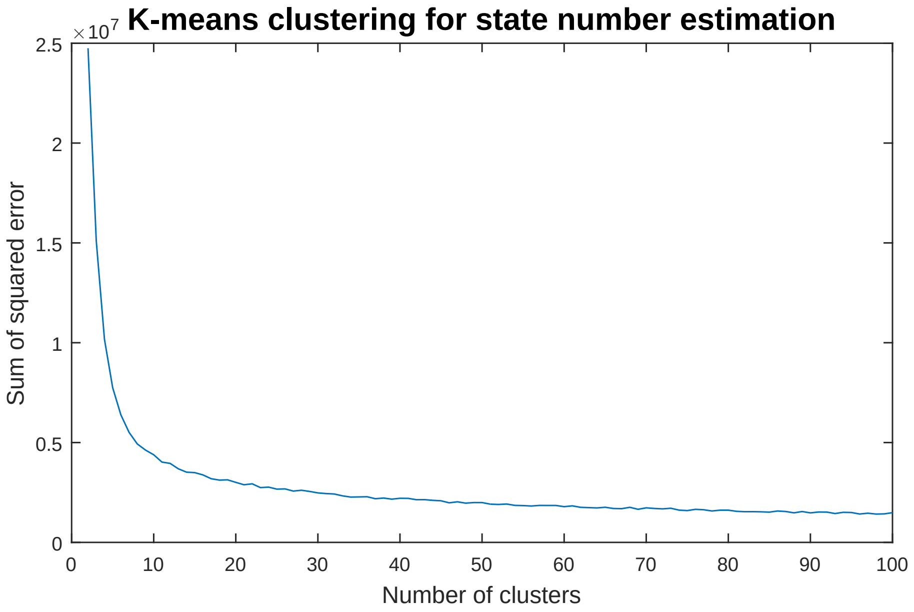


**Figure S3:** The change of Sum of Squared Error (SSE) with respect to an increased number of clusters in $K$-means clustering on the cross-species Hi-C data. The number of hidden states was estimated to be between 25 and 35 based on the results from $K$-means clustering.

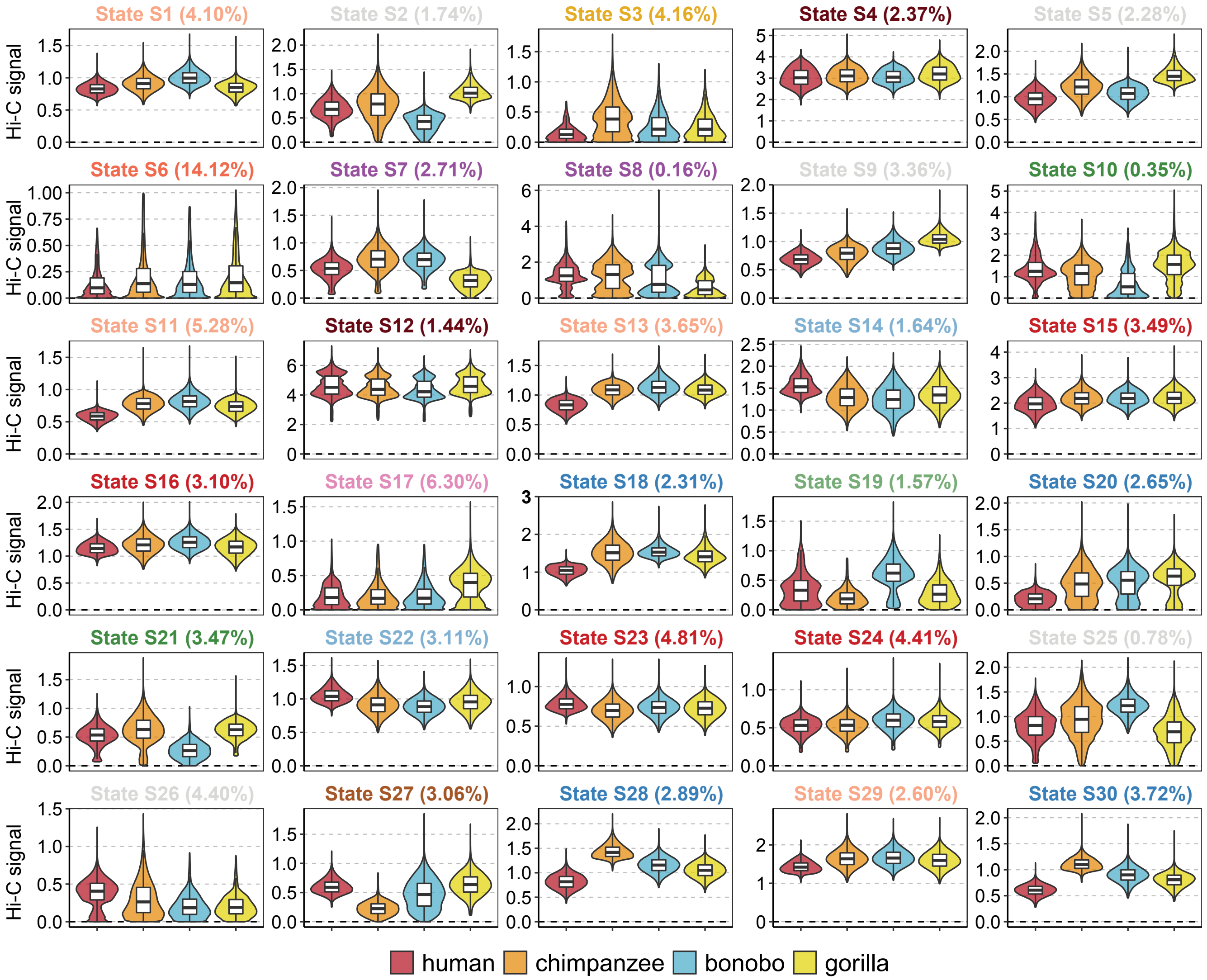


**Figure S4:** Hi-C evolutionary patterns identified by Phylo-HMRF in all the major synteny blocks on all autosomes across four primate species. The boxplot shows the normalized cross-species Hi-C contact frequency distributions in the corresponding state. The color of the boxplot title shows the evolutionary group assignment of the corresponding state.

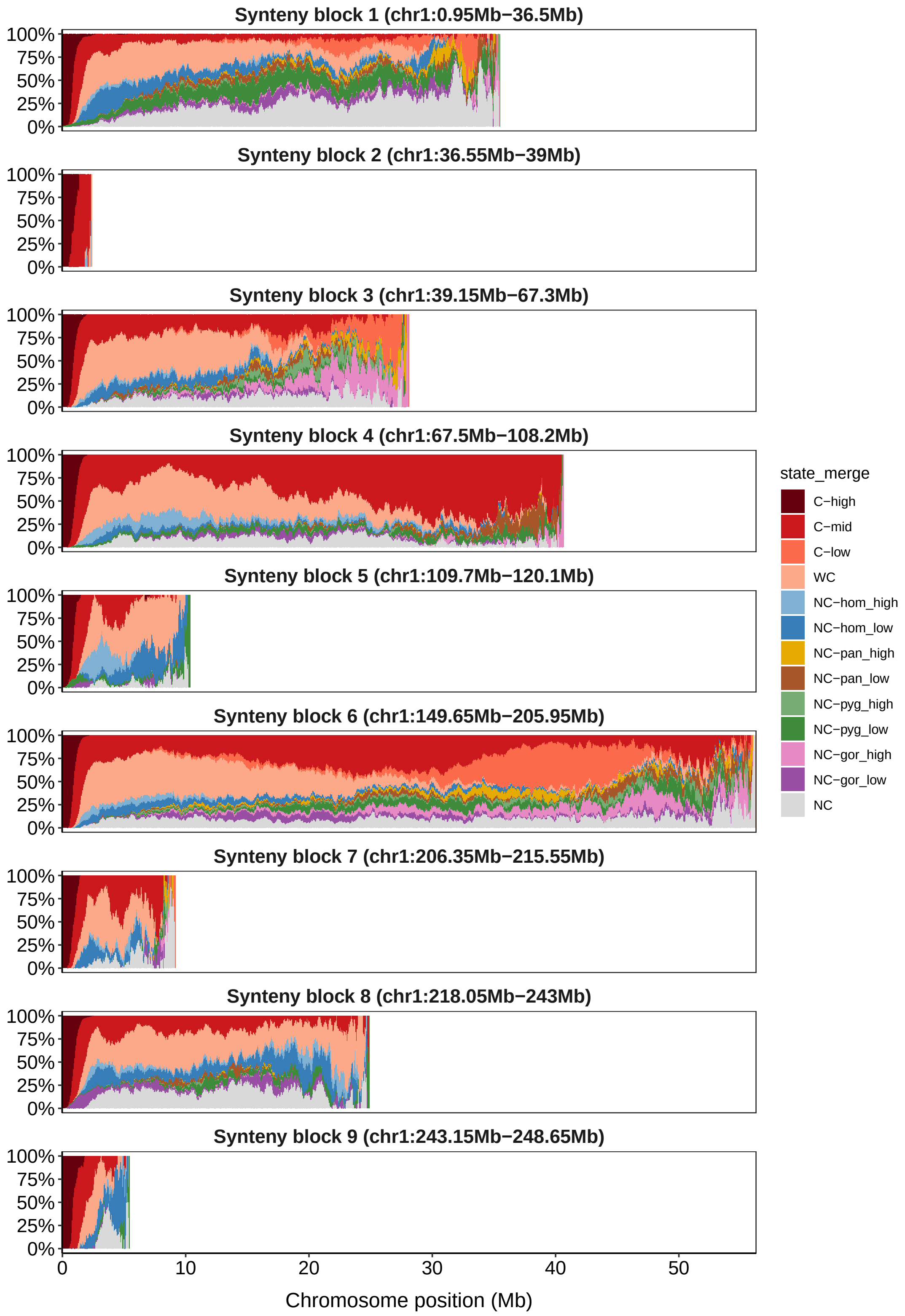


**Figure S5:** Distributions of predicted Hi-C contact evolutionary states over changing distance between a pair of genomic loci in each synteny block on human chromosome 1.

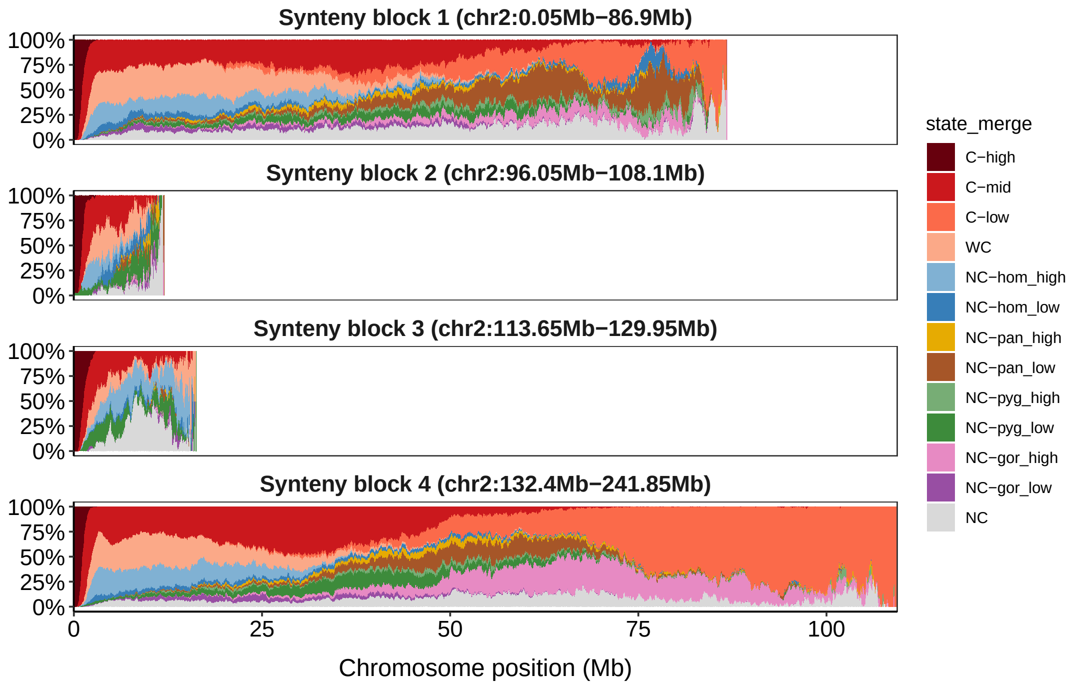


**Figure S6:** Distributions of predicted Hi-C contact evolutionary states over changing distance between a pair of genomic loci in each synteny block on human chromosome 2.

| Simulation | Method | NMI | AMI | ARI | Precision | Recall | $F_1$ |
|---|---|---|---|---|---|---|---|
| Dataset I-1 | Clustering | 0.1096 | 0.1001 | 0.0532 | 0.2445 | 0.1384 | 0.1767 |
| Dataset I-1 | GMM | 0.1213 | 0.1115 | 0.0603 | 0.2502 | 0.1497 | 0.1873 |
| Dataset I-1 | SLIC | 0.0602 | 0.0554 | 0.0272 | 0.2117 | 0.1280 | 0.1595 |
| Dataset I-1 | Quickshift | 0.033 | 0.0281 | 0.0265 | 0.1972 | 0.3173 | 0.2433 |
| Dataset I-1 | Gaussian-HMRF | 0.6214 | 0.6046 | 0.6981 | 0.8068 | 0.6985 | 0.7487 |
| Dataset I-1 | Phylo-HMRF | 0.6957 | 0.6915 | 0.7786 | 0.8373 | 0.7989 | 0.8177 |
| Dataset I-2 | Clustering | 0.1457 | 0.1074 | 0.0520 | 0.5626 | 0.1349 | 0.2176 |
| Dataset I-2 | GMM | 0.1936 | 0.1434 | 0.0793 | 0.6118 | 0.155 | 0.2474 |
| Dataset I-2 | SLIC | 0.0776 | 0.0573 | 0.0042 | 0.4604 | 0.1113 | 0.1793 |
| Dataset I-2 | Quickshift | 0.0519 | 0.0412 | 0.0002 | 0.4519 | 0.1579 | 0.2341 |
| Dataset I-2 | Gaussian-HMRF | 0.4279 | 0.3368 | 0.3150 | 0.8547 | 0.3470 | 0.4933 |
| Dataset I-2 | Phylo-HMRF | 0.6346 | 0.6191 | 0.8293 | 0.9293 | 0.8807 | 0.9043 |
| Dataset I-3 | Clustering | 0.2046 | 0.1900 | 0.1269 | 0.2938 | 0.1956 | 0.2348 |
| Dataset I-3 | GMM | 0.2306 | 0.2183 | 0.1728 | 0.3237 | 0.2558 | 0.2857 |
| Dataset I-3 | SLIC | 0.0724 | 0.0676 | 0.0318 | 0.1881 | 0.1329 | 0.1557 |
| Dataset I-3 | Quickshift | 0.0595 | 0.0500 | 0.0198 | 0.1658 | 0.3791 | 0.2307 |
| Dataset I-3 | Gaussian-HMRF | 0.6447 | 0.6304 | 0.6854 | 0.7534 | 0.7131 | 0.7327 |
| Dataset I-3 | Phylo-HMRF | 0.7344 | 0.7327 | 0.8010 | 0.8288 | 0.8350 | 0.8319 |
| Dataset I-4 | Clustering | 0.1003 | 0.0850 | 0.0332 | 0.3526 | 0.1207 | 0.1798 |
| Dataset I-4 | GMM | 0.1191 | 0.1017 | 0.0455 | 0.3685 | 0.1355 | 0.1982 |
| Dataset I-4 | SLIC | 0.0705 | 0.0602 | 0.0159 | 0.3224 | 0.1183 | 0.1731 |
| Dataset I-4 | Quickshift | 0.044 | 0.0437 | 0.0054 | 0.3025 | 0.2095 | 0.2475 |
| Dataset I-4 | Gaussian-HMRF | 0.3699 | 0.3179 | 0.1921 | 0.5872 | 0.2249 | 0.3252 |
| Dataset I-4 | Phylo-HMRF | 0.5480 | 0.5275 | 0.6253 | 0.8297 | 0.6396 | 0.7223 |
| Dataset I-5 | Clustering | 0.1429 | 0.1368 | 0.0657 | 0.2061 | 0.1594 | 0.1798 |
| Dataset I-5 | GMM | 0.1561 | 0.1508 | 0.0737 | 0.2100 | 0.1782 | 0.1928 |
| Dataset I-5 | SLIC | 0.0620 | 0.0593 | 0.0244 | 0.1644 | 0.1282 | 0.1441 |
| Dataset I-5 | Quickshift | 0.0295 | 0.0264 | 0.0073 | 0.1451 | 0.2138 | 0.1729 |
| Dataset I-5 | Gaussian-HMRF | 0.4990 | 0.4833 | 0.4776 | 0.5968 | 0.5004 | 0.5443 |
| Dataset I-5 | Phylo-HMRF | 0.6129 | 0.6074 | 0.6442 | 0.7160 | 0.6706 | 0.6924 |
| Dataset I-6 | Clustering | 0.1327 | 0.1137 | 0.0760 | 0.3647 | 0.1561 | 0.2186 |
| Dataset I-6 | GMM | 0.1697 | 0.1479 | 0.1087 | 0.3944 | 0.1939 | 0.2600 |
| Dataset I-6 | SLIC | 0.0574 | 0.0491 | 0.0093 | 0.2707 | 0.1148 | 0.1613 |
| Dataset I-6 | Quickshift | 0.0502 | 0.0488 | 0.0233 | 0.2770 | 0.2318 | 0.2524 |
| Dataset I-6 | Gaussian-HMRF | 0.4837 | 0.4572 | 0.6622 | 0.8029 | 0.6907 | 0.7426 |
| Dataset I-6 | Phylo-HMRF | 0.6203 | 0.6067 | 0.7580 | 0.8582 | 0.7802 | 0.8173 |
| Dataset I-7 | Clustering | 0.1547 | 0.1379 | 0.0926 | 0.3506 | 0.1732 | 0.2318 |
| Dataset I-7 | GMM | 0.2250 | 0.2013 | 0.1874 | 0.4632 | 0.2397 | 0.3157 |
| Dataset I-7 | SLIC | 0.1066 | 0.0946 | 0.0370 | 0.2809 | 0.1334 | 0.1809 |
| Dataset I-7 | Quickshift | 0.0577 | 0.0502 | 0.0191 | 0.2451 | 0.2965 | 0.2684 |
| Dataset I-7 | Gaussian-HMRF | 0.5072 | 0.4754 | 0.5452 | 0.7254 | 0.5694 | 0.6380 |
| Dataset I-7 | Phylo-HMRF | 0.5506 | 0.5190 | 0.5819 | 0.7561 | 0.5975 | 0.6675 |
| Dataset I-8 | Clustering | 0.1019 | 0.0828 | 0.0153 | 0.3396 | 0.1165 | 0.1735 |
| Dataset I-8 | GMM | 0.1281 | 0.1052 | 0.0246 | 0.3514 | 0.1333 | 0.1933 |
| Dataset I-8 | SLIC | 0.0873 | 0.0710 | 0.0255 | 0.3558 | 0.1236 | 0.1834 |
| Dataset I-8 | Quickshift | 0.0557 | 0.0553 | 0.0457 | 0.3513 | 0.2835 | 0.3138 |
| Dataset I-8 | Gaussian-HMRF | 0.3286 | 0.2746 | 0.1733 | 0.5354 | 0.2457 | 0.3368 |
| Dataset I-8 | Phylo-HMRF | 0.5897 | 0.5618 | 0.6550 | 0.8405 | 0.6805 | 0.7520 |

**Table S1:** Performance evaluation of $K$-means Clustering, GMM, SLIC, Quick Shift, Guassian-HMRF, and Phylo-HMRF on eight simulated datasets in Simulation Study I using evaluation measurements NMI, AMI, ARI, Precision, Recall, and $F_1$ score. Each method is repeated 10 times on each simulation dataset with different initializations if applicable. The average performance from the 10 repeated runs of each method is presented.
